# Estimation, testing, and inference of network heterogeneity

**DOI:** 10.64898/2025.12.18.695221

**Authors:** Zhanshan (Sam) Ma, Aaron M. Ellison

## Abstract

1. Diversity and heterogeneity are related but distinct and often conflated concepts. Diversity quantifies the number or relative abundance of discrete objects (e.g. species), whereas heterogeneity includes interactions among them (i.e. in networks) and between them and their environments. Although estimation, testing, and inference of diversity is well established and understood in ecology, comparable methods for heterogeneity are themselves diverse and rarely applied consistently or coherently.
2. We propose a consistent and coherent methodology for estimation, testing, and inference of heterogeneity of ecological networks. Estimation of heterogeneity is scalable from individuals to populations using the variance-to-mean (*V*/*M*) ratio and extensions of Taylor’s power law (TPL) to analyzing networks. Bootstrapping is used to partition heterogeneous and random clusters, whereas permutation tests are used to compare individual- and network-level heterogeneity. Inference includes the identification of “important” (e.g. dominant, foundation, keystone) species and “rich clubs” in heterogeneous networks, detection of biomarkers, and analysis of heterogeneity–stability relationships.
3. We demonstrate this methodology using the global Earth Microbiome Project dataset. The method could reliably distinguish heterogeneous nodes and networks; identified significant differences in heterogeneity among microbial assemblages in different habitats and in specific sites within habitats; and supported established principles of host filtering, species sorting, and niche partitioning.
4. Our methods for estimation, testing, and inference of heterogeneity are modular, scalable, and applicable to a wide range of ecological systems. They also provide a quantitative method for understanding how evolutionary and ecological forces jointly shape both topology and heterogeneity in ecological networks.

## INTRODUCTION

Diversity and heterogeneity are related but distinct and often conflated concepts. Shavit & Ellison (2021) suggested an aphorism—”a zoo is diverse, an ecosystem is heterogeneous”—to encapsulate the distinction between them. That is, diversity quantifies the variety (e.g. number, relative abundance) of discrete objects (e.g. species), whereas heterogeneity includes interactions between them (Shavit & Ellison, 2021; see also Ma, 2025). Ecologists describe and analyze diversity with well-understood measures (e.g. Hill numbers; Chao et al., 2014; and papers in Ellison, 2010), but a comparable set of measures and methods for quantifying and analyzing heterogeneity (sensu Shavit & Ellison, 2021) remains underdeveloped.

The lack of a clear and consistent way of quantifying heterogeneity stems from a lack of an agreed-upon definition of it (e.g. Kolasa & Rollo, 1991; Sparrow et al., 1999; Li & Reynolds, 1995; Stein & Kreft, 2015; Ma & Ellison, 2026). For example, Kolasa & Rollo (1989) tried to match different definitions of heterogeneity to different systems, scales, and questions. Dutilleul & Legendre (1993) differentiated ecological heterogeneity from the purely statistical concept of heteroscedasticity, whereas Li & Reynolds (1995) distinguished different measures of heterogeneity for different kinds of data. Heterogeneity also has been defined differently for different spatial and temporal scales and levels of biological organization (e.g. Cadenasso et al., 2006; Vinatier et al., 2010; Allouche et al., 2012; Collins et al., 2018; Loke & Chisholm, 2022; Ma & Ellison, 2026).

For any type of ecological data and spatiotemporal scale, however accounting for interactions is a necessary condition for any measure of heterogeneity (Shavit & Ellison, 2021; Ma, 2025). Because specifying, measuring, and analyzing interactions is straightforward for networks of interacting species (such as bipartite networks of plants and pollinators or food webs), they should be a good starting point from which to develop a coherent methodology for measuring and estimating heterogeneity. Indeed, Forsythe et al. (2021) applied Estrada’s (2010) general method for quantifying network heterogeneity to model how among-individual variation (“individual heterogeneity”) in vital rates scale up to influence population dynamics. Zelnik et al. (2023) introduced the idea of “ecological collectivity” that has much in common with Shavit & Ellison’s (2021) concept of heterogeneity. Zelnik et al. (2023) measured ecological collectivity as the spectral radius of a community interaction matrix and also quantified the extent of indirect effects in the community.

Here, we use concepts from network science and a graph-based formalism to develop a general, scalable method for the estimation, testing, and inference of ecological heterogeneity (sensu Shavit & Ellison, 2021). We then illustrate this method by applying it to the Earth Microbiome Project (EMP) dataset (Thompson et al., 2017). Our approach is inspired by the hierarchical, modular, and robust design of the Open Systems Interconnection (OSI) reference model (ISO/IEC 7498-1; Zimmermann, 1980). The OSI model underpins all digital networks, including the Internet and wireless networks, and includes hierarchical, modular, scalable, and interoperable protocol stacks that ensure reliable communication between network nodes. We do not mean to imply that ecological networks function like the Internet. Rather, the hierarchical complexity, scalability, and interoperability of computer communication networks inspired our thinking about developing general and robust methods for estimating heterogeneity in complex ecological networks.

### DEFINING, MEASURING, AND TESTING NETWORK HETEROGENEITY

We begin with definitions and measurements of node and network heterogeneity (summarized in Table 1), and then proceed to parameter estimation, testing, and inference. Throughout, we consider networks whose nodes (species) and edges (interactions) are “weighted” by, respectively, their abundances and interaction strengths (e.g. Emmerson et al., 2005). We emphasize that we want to distinguish how individual species with varying abundances interact with other species in the network. We also want to understand scaling relationships between the heterogeneities of parts of a network (e.g. its nodes, edges, or subwebs) and the heterogeneity of the entire network.

**Table 1.** Summary of key definitions and formulas to compute node and network heterogeneity.

| Definition | Formula | Interpretation |
| --- | --- | --- |
| Node heterogeneity ($H_i$; Eqn. 4). | $H_i = \frac{V_i}{M_i}$ | $V_i$ and $M_i$ are the variance and mean of the elements of the vector of *weighted species connectedness* $C_i^W$ (Eqn. 3). |
| Heterogeneity of fully weighted networks (${}^qH$; Eqns. 5, 6). | ${}^qH = \left( \sum_{i=1}^S h_i^q \right)^{1/(1-q)}$ | $q$ is the Hill number exponent and $h_i = \frac{H_i}{\sum_{i=1}^S H_i}$ is the relative heterogeneity of node $i$. |
| Power-law probability distribution for scaling and heterogeneity modeling (Eqn. 9) | $P(x) = \frac{\alpha - 1}{x_{min}} \left( \frac{x}{x_{min}} \right)^{-\alpha} \propto (x)^{-\alpha}$ | A smaller $\alpha$ (e.g. $1 < \alpha < 2$) indicates heavier tail or high heterogeneity in species interactions. When $x > x_{min}$, power-law behavior emerges. |
| Taylor's Power Law for scaling network heterogeneity (Eqn. 9; after Ma [2025]). | $V = aM^b$ | $V$ and $M$ are all the pairs of $\{V_i, M_i\}$ computed from $C_i^W$ (Eqn. 3). |

Paying attention to differences in both numbers and strengths of interactions between species with different abundances differs substantially from how network ecologists normally measure topological properties of unweighted networks such as food webs (e.g. Dunne et al., 2002; McCann, 2011). A familiar example is connectance, 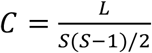, where *S* is the number of species in the web (without information on their relative abundances or biomass) and *L* is the total number of links between them (without information on their interaction strengths).

In contrast, we start by defining the *connectedness* of an individual species *i* in a network as:

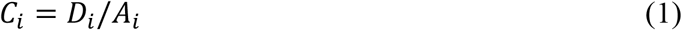

where *D*_*i*_ is the node-degree of species *i* (that is, the number of links to or from *i*) and *A*_*i*_ is the mean abundance of species *i* across all the samples used to construct the network (note that our Eqn. 1 is the same as Eqn. 3 in Ma [2025], who equated *C*_*i*_ with the number of significant correlations between species *i* and all other species in a microbial network). To account for interaction strength, we replace *D*_*i*_ with its link-weighted sum (*LWSum*)

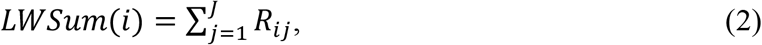

where *R*_*ij*_ is the strength of the interaction between species *i* and *j* for the *J* species that share links with species *i*. With this replacement, species connectedness is 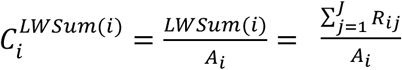. For weighted networks, 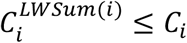 (as defined in Eqn. 1), because |*R*_{*ij*}_ ≤ 1| and 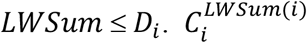 also is equivalent to the node-strength metric that Bertolero et al. (2017) used to identify rich clubs in complex networks.

Species connectedness as defined in Eqn. (1) is simple and relatively coarse, whereas the alternative link-weighted sum version (Eqn. 2) appears more fine-grained since it sums up actual weight values rather than binary presence/absence of links. However, we found in previous work (Ma, 2025) and in our preliminary explorations with the EMP dataset used herein that the additional finer-grained information of 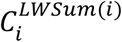 failed to deliver substantially better or more stable estimates of connectedness.

Ma (2025) suggested that a more effective way to use link weights would be to compute the *weighted species connectedness* of species *i*:

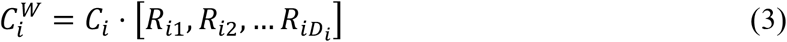

where [*R*_*ij*_] represents the vector of interaction strengths between species *i* and *j* ∈{1, 2, …, *D*_*i*_} and *D*_*i*_ is the node-degree of species *i*. Note that 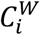 is not a scalar, but rather is the scalar product (and hence a vector) of the product of the species connectedness (the scalar, *C*_*i*_, from Eqn. 1) and the vector of the weights of the direct links between nodes (interaction strengths, correlation coefficients, etc.).

#### Heterogeneity of individual nodes

Using Eqns. 1 & 3, we define the *node heterogeneity* (*H*_*i*_) for each node *i* using the mean *M*_*i*_ and variance *V*_*i*_ of the vector 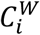 as:

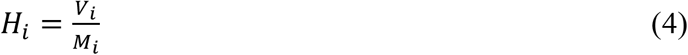

with *H*_*i*_ ∈ [0, ∞) . Eqn. 4 assumes that each species (node) exhibits intrinsic heterogeneity through aggregated variation in three key attributes: (1) abundances (node weights), (2) connectivity (node degree), and (3) interaction strengths (edge weights). Node heterogeneity thus quantifies individual dispersion from mean connectedness while also incorporating the topological influences of neighboring nodes through their patterns of connectivity.

To account for noise in estimates of *H*_i_, we used a bootstrap test to filter artifacts and estimate node heterogeneity, and to partition nodes into clusters based on tests for whether *H*_i_ > 1 (heterogeneous) and another for *H*_*i*_ < 1 (uniform). Nodes failing both tests are classified as random. The computational algorithm for doing these bootstrap tests is summarized in Appendix A.

#### Heterogeneity of weighted networks

We define *network heterogeneity* by aggregating node heterogeneities using Hill numbers (Rényi entropy):

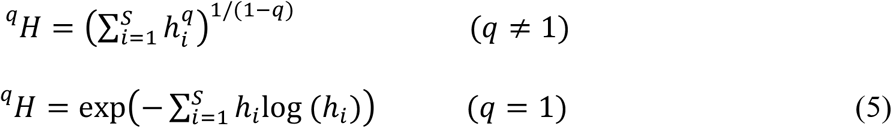

where *h*_*i*_ is the *relative node heterogeneity*:

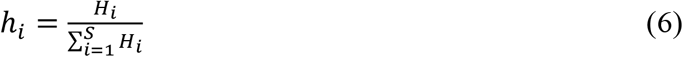

and *H*_*i*_ is the node heterogeneity (Eqn. 4). In parallel with Hill numbers for measuring diversity, ^0^*H* is simply the species richness or the number of network nodes, ^1^*H* is the number of equivalent species with identical node heterogeneities, and ^2^*H* is the number of equivalent species with dominant node heterogeneities.

In the absence of link weights, we could substitute *relative node abundance* 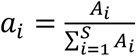 for *h*_*i*_, where *S* is the number of species in the network and *A*_*i*_ is the mean abundance of species *i* across all samples used to build the network. This substitution simply represents an extension of Hill number diversity indices to networks. Similarly, in the absence of node weights, we could substitute *relative link weight* 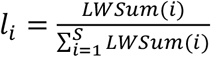, where *LWSum*(*i*) was defined in Eqn. 2. This substitution gives more information about species interactions, but as it lacks node (species abundance) weights, it is, at best, only a minimal measure of network heterogeneity.

To characterize the change in network heterogeneity with increasing values of *q*, we define a scaling exponent λ as:

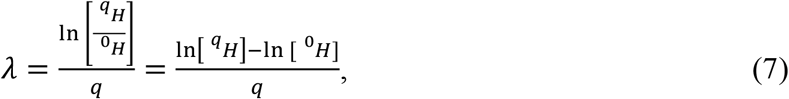

with ^*q*^*H* as in Eqn. 5. Because ^*q*^*H* declines with *q, λ* < 0. More negative λ values indicate greater heterogeneity for fully-weighted networks, stronger dominance effects for species abundance in node-weighted networks, and greater inequality for interaction strengths in link-weighted networks.

Alternatively, we could define network heterogeneity as 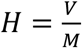, where *V* = var (*H*_*i*_) and *M* = var (*H*_*i*_); *H*_*i*_ as in Eqn. 4. In this formulation, *H* measures statistical dispersion at the network level: *H* = 1 implies a randomly structured network; *H* > 1 implies a heterogeneous network; and *H* < 1 a (more) uniform network. Because quantifying network heterogeneity in this way is overly simplistic, we think it better to model it probabilistically from power-law distributions of node heterogeneity (e.g. Clauset et al., 2009).

The power-law probability density function is

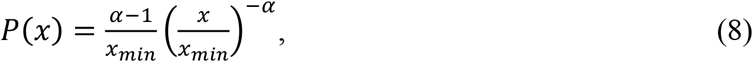

where *x*_min_ and *α* are threshold and scaling parameters, respectively. The key feature of Eqn. 8 is that *P*(*x*) ∝ (*x*)^−*α*^; smaller *α* values imply a heavy-tailed (more slowly-decaying) distribution. In network science, heavy tails indicate a few extreme hubs (scale-free networks), whereas in ecology, heavy tails indicate high heterogeneity in species interactions. When *x < x*_*min*_, *p*(*x*) = 0, and the data may be fit better by an exponential or uniform distribution. Hence, *x*_*min*_ is a minimum (threshold) value above which power-law behavior holds. Technical details for estimating the scaling exponent α and the parameter *x*_*min*_ are given in Appendix B.

The scaling exponent *α* also determines key statistical and ecological properties of power-law distributions. For 1 < *α* ≤ 2, *P*(*x*) has infinite mean and variance, reflecting extreme heterogeneity and frequent rare events characteristic of systems with dominant hub-like elements. When 2 < *α* ≤ 3, the mean is finite but the variance remains infinite, indicating substantial but bounded fluctuations in heterogeneity, as often seen in metacommunity dynamics. Distributions with *α* > 3 have finite mean and variance, corresponding to lighter-tailed distributions and reduced ecological heterogeneity typical of more stable systems. Each of these distributions also implies distinct mechanisms governing species interactions and abundance patterns.

Although the power-law distribution has been used previously to measure network heterogeneity (Ma, 2025), the innovation here is in using it to fit the distribution of node heterogeneity (via *V*_*i*_/*M*_*i*_) rather than directly fitting node degree or abundance distributions (Appendix B). To compare heterogeneities between networks, we can use *Z*-tests (for comparisons of the scaling exponent α) and Kolmogorov-Smirnov tests (to compare full distributions). Technical details of these methods are given in Appendix C.

Finally, we can also scale heterogeneity by applying Taylor’s Power Law of Networks (TPLoN; Ma, 2025) to the {*V*_*i*_, *M*_*i*_} pairs (Eqn. 4) computed from each 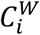 (Eqn. 3) across all the network nodes:

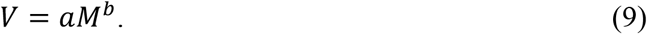

We then use permutation tests on node heterogeneity, network heterogeneity, and TPLoN network heterogeneity scaling parameters to identify nodes with significant changes in heterogeneity, including nodes that switch between clusters and those that change their heterogeneity but remain within a particular cluster, and to detect global differences in heterogeneity between two networks (Appendix D).

### INFERENCE

#### Heterogeneity of nodes

The bootstrap tests (Appendix A) group nodes into heterogeneous, random, and uniform clusters. Because it is based on statistically defined properties of nodes, our approach differs from existing methods such as core/periphery partitioning (Gallagher et al., 2021) and molecular complex detection (MCODE; Bader & Hogue, 2003). We also can use our method to infer different types of “clubs” in networks (Bertolero et al., 2017). Specifically, we define and identify rich clubs and heterogeneous clubs.

For a network with up to three clusters of nodes (heterogeneous, random, and uniform), we define the participation coefficient (PC) of node *i* as:

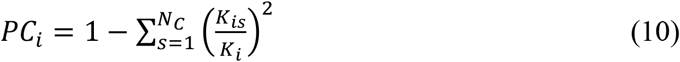

where *K*_*i*_ is the sum of node *i*’s edge weights (equal to the link-weighted sum, *LWSum*(*i*); Eqn. 2), *K*_*is*_ is the sum of node *i*’s edge weights to a cluster *S*, and *N*_*C*_ is the total number of clusters in the network (1 ≤ *N*_*C*_ ≤ 3). *PC* measures how evenly distributed a node’s edges are across the clusters. *PC* is maximized when it has an equal sum of edge weights in each cluster in the network. *PC* = 0 if all edges are to a single cluster. Note that *PC* resembles Simpson’s diversity index and analogously quantifies node heterogeneity by measuring the unevenness of connections across network modules. Both *PC* and Simpson’s index are sensitive to the skewness of their respective components (respectively, link weights and species abundances).

Operationally, heterogeneous club nodes are defined by those nodes whose *PC* ≥ the 80^th^ percentile of measurements of cross-module connectivity between node clusters. Rich-club nodes are nodes for which *K*_*i*_ ≥ the 80^th^ percentile of all node-strengths *K*_*i*_. In general, rich clubs are dominated by connections within clusters whereas heterogeneous clubs bridge modules and are dominated by nodes with high participation coefficients.

#### Network heterogeneity

Our measure of heterogeneity of fully weighted networks (Eqns. 5, 6) brings together measurements of both node-level and link-level heterogeneity. It is essential to distinguish between the magnitude of heterogeneity at a given scale from how heterogeneity changes across scales (see previous subsections and Appendices B–D). The scaling parameters in Eqns. 8 and 9 (respectively, α and *b*) clarify this difference. We can also use TPLoN to infer differences in magnitude of network heterogeneity (parameter *b* in Eqn. 9; Ma, 2025).

In addition, the fitted parameters *a* and *b* in Eqn. 9 can be used to estimate *M*_0_, which Ma (2015, 2025) called the community heterogeneity criticality threshold (HCT):

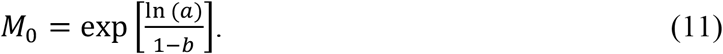

*M*_0_ also is equal to the mean weighted connectedness for which *V*/*M* = 1 (random heterogeneity). When *M* in Eqn. 9 = *M*_0_, we infer that external drivers are unimportant in determining network heterogeneity (analogous to Eisler et al.’s [2008] illustration of a system’s internal fluctuations driving its endogenous behavior). In contrast, when *M* > *M*_0_, external drivers are inferred to be significant drivers of network heterogeneity; when *M* < *M*_0_, we infer that networks are more uniform (homogeneous) than random.

### CASE STUDY: THE EARTH MICROBIOME PROJECT (EMP) DATASET

#### The dataset

The Earth Microbiome Project (EMP) includes 23,727 samples from host-associated (*N*=11,358) and free-living (*N*=12,369) microbial communities (Table S1; Thompson et al., 2017). The EMP has been used extensively to study ecological patterns and processes in microbial ecology ranging from small-scale host-microbe interactions to large-scale biogeographic patterns on land, in soils, and in the oceans.

The value of the EMP’s dataset as a case-study for examining heterogeneity among microbial networks derives from its standardized metagenomic processing method across ecosystems that allows for construction and comparison of weighted interaction networks. The EMP dataset includes 15 “treatments”, which we aggregated into six broader “groups”: animal, plant, sediment, soil, surface, and aquatics, each of which may be divided into finer “sites” within groups (Table S1). To further assess differences in microbial community heterogeneity across environmental (habitat) and host-associated gradients, we also defined two additional pairs of groups: non-saline vs. saline and free-living vs. host-associated. In total, this yielded 10 groups and 14 group × site combinations for subsequent analyses of heterogeneity across hierarchical scales (Table S1).

The data from each group or group × site combination (Table S1) was then formatted as an *m × n* (“site × species”) matrix, with *m* rows of biological samples and *n* columns of microbial OTUs; each entry {*m*_*i*_, *n*_*j*_} in the matrix is the abundance of OTU *j* in sample *i*. To minimize the effects of spurious reads, we removed OTUs (Operational Taxonomic Units) with fewer than 10 reads or that occurred in < 5% of samples in each of the 24 group or group × site combinations. We then constructed ecological correlation networks (one for each group or group × site combination) using the Weighted Network (WN) algorithm using the WGCNA package (Langfelder & Horvath 2008). The 24 WN datasets are in Datasets S1 & S2; metadata for them are in Tables S1 & S2). The WN approach is preferred over alternatives such as the SparCC algorithm (Friedman & Alm, 2012) because WN preserves all possible links by standardizing the link weight assignment with transformation algorithms that filter out noise, and it standardizes all weights to the interval [0, 1] (Langfelder & Horvath 2008).

#### Node heterogeneity, network clusters, and heterogeneous clubs

Summary statistics for each OTU—mean node abundance, link-weighted sums, node heterogeneities, participation coefficients, and assignments to heterogeneous, uniform, or random clusters (based on bootstrap tests), rich clubs (based on link-weighted sums), or heterogeneous clubs (based on participation coefficients)—in each of the WNs are given in Datasets S1 and S2. Summaries of overall network characteristics, including raw and bootstrap-validated percentages of heterogeneous, random, and uniform nodes, and raw and bootstrapped mean node heterogeneity values are in Table S4. The bootstrap-validated values of each of these allow us to distinguish true node heterogeneity from artifacts detected using only *V*/*M*. Five important findings or patterns emerged from these analyses (summarized in Table S4; Fig. 1 illustrates these for a single example: the animal [group] × corpus [site] microbiome network):

1. Heterogeneity was present at both node (OTU) and network levels. The mean node heterogeneity consistently exceeded 1 across all groups and group × site combinations.
2. Bootstrap testing reduced the estimates of the number of heterogeneous and uniform nodes, and increased the estimated number of random nodes.
3. Heterogeneity of nodes differed among groups (habitats). Specifically,
  a. sediments exhibited extreme heterogeneity (95–96% bootstrap-validated);
  b. aquatic groups showed moderate heterogeneity (53–75% bootstrap-validated);
  c. host-associated microbiomes had somewhat lower heterogeneity than free-living microbial assemblages (∼88% vs. 93% bootstrap-validated).
4. Rich-club members, which exhibit high within-module connectivity, function as key local connectors in their respective modules. For example, the heterogeneous species cluster in saline sediments combined extreme node heterogeneity (*V*/*M*=219.838) with a high link-weighted sum (*K*_*i*_ = 63.085).
5. Participation coefficients (*PC*) revealed distinct connectivity patterns: the heterogeneous species cluster in the animal corpus showed high cross-module connectivity (*PC*=0.363; Fig. 1), whereas the heterogeneous species cluster in the plant rhizosphere exhibited low cross-module connectivity (PC=0.021).

The power-law distribution was the only distribution that fit at least some of the observed data (42% fit rates; Table S5); excellent fits were observed for microbiomes of the animal corpus (*P* = 0.542), plant corpus (*P* = 0.950), and grouped animal samples (*P* = 0.694). The poorest power-law fits consistently occurred in surface-associated habitats across multiple groups. Scaling exponents (α) of the fitted power-law distributions ranged from 1.649 to 4.118, and minimum threshold values (*x*_min_) spanned nearly two orders of magnitude (3.3–208.5). Host-associated samples exhibited the steepest scaling (*α* = 4.118), surface habitats consistently showed lower (shallower) scaling than other groups or habitats. Finally, nearly all between-group or between group × site combinations exhibited significant differences in scaling parameters (Tables S6, S7).

**Figure 1.**
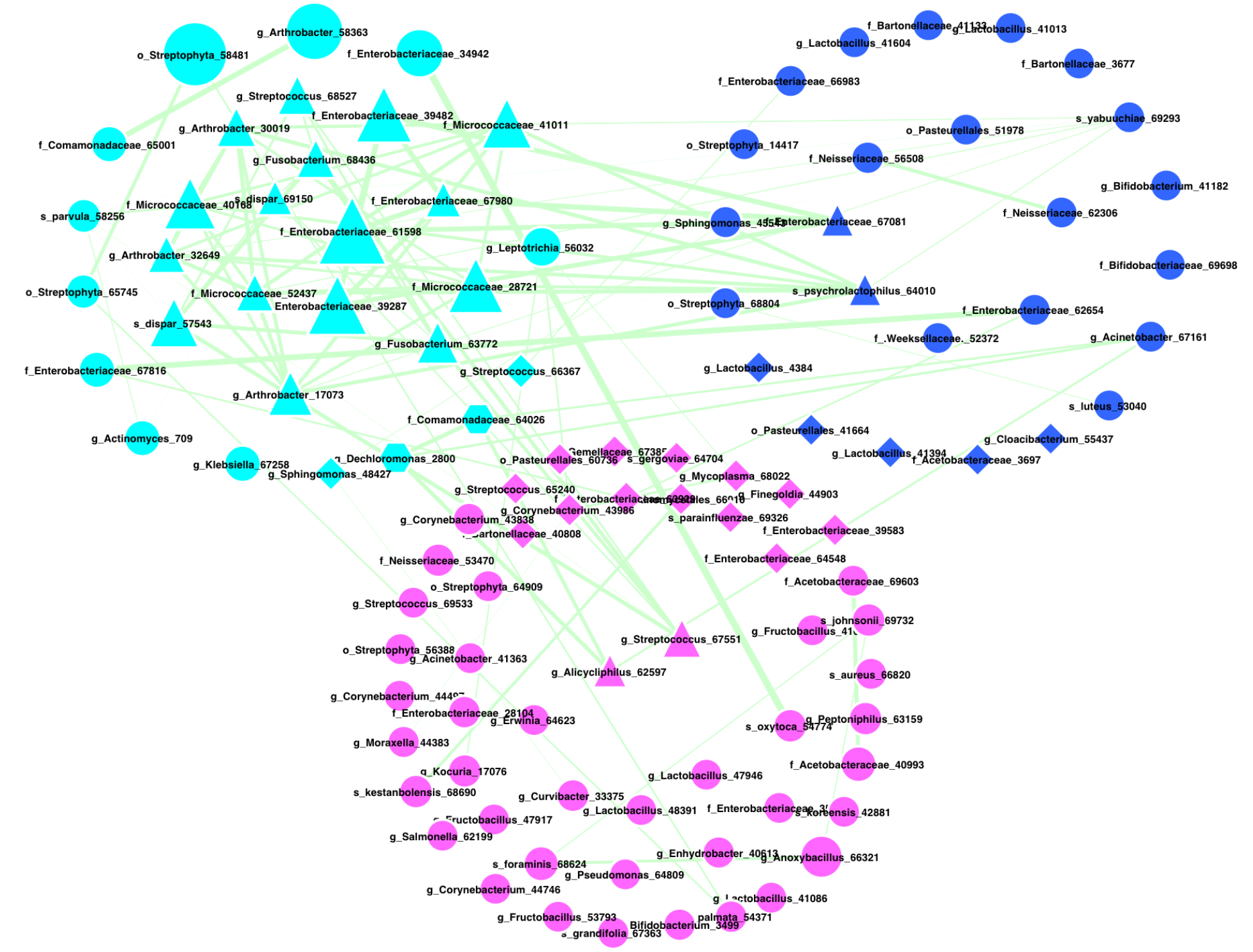
Partitioning of the animal corpus microbiome network into three node clusters—heterogeneous (cyan), random (blue), and uniform (magenta)—based on bootstrap testing of node heterogeneity. Rich club nodes are shown as triangles, heterogeneous club nodes as diamonds, and dual-role nodes (rich club + heterogeneous club) as hexagons. The magnitude of node heterogeneity *H*_*i*_ is reflected in the size of the symbol and link thickness represents estimated weight of the edges.

#### Network heterogeneity with varying Hill numbers

The absolute value of the scaling exponent λ (Eqn. 7) reflects differences (decay) in strength as the Hill exponent *q* changes. Larger values of |λ| values indicate sharper species dominance in node abundance (i.e. classical biodiversity as measured by Hill numbers), more disproportionate connections in link weights, and stronger imbalance in overall network heterogeneity. These three measures of network heterogeneity exhibited distinct scaling patterns in the EMP (Table S8):

1. *Network heterogeneity of node abundance* had the steepest decays (λ = −0.76 – −0.15), indicating high species dominance (extreme unevenness in abundances) in which a few dominant species determine system-wide heterogeneity in species abundances.
2. *Network heterogeneity of link weights* decayed less steeply (λ = −0.35 – −0.08). This result suggests that evolutionary constraints maintain relatively equitable link-weight distributions despite differences in node abundances (Ma & Ellison, 2024).
3. *Overall network heterogeneity*, which incorporates both node and edge heterogeneity, exhibited intermediate scaling behavior (λ = −0.55 – −0.22) that directly reflects the competing influences of evolutionary constraints in host-associated systems and ecological opportunity in free-living environments. Specifically:
  a. Host environments impose evolutionary filters that systematically reduce heterogeneity, as evidenced by characteristically shallow scaling (λ ≈ −0.21) that likely arises from physiological selection preferentially favoring dominant taxa; niche compression limiting rare-species persistence; and coevolution stabilizing interactions. These forces collectively reduce network topology and constrain heterogeneity, resulting in 60% shallower scaling in host-associated microbiomes than in free-living systems, 2.1× lower rare-species contributions (*q*=0→1), and 45% stronger dominance effects.
  b. In contrast, free-living microbial assemblages had steeper scaling (λ = −0.55 – −0.50. Non-saline sediment assemblages exhibited the strongest rare-species contributions (λ = −0.55), followed by non-saline waters (λ = −0.50) and soil habitats (λ = −0.52).

We used permutation tests to test for differences in the three types of network heterogeneity (of node abundance, link-weight, and overall) across different environments (microbial habitats and hosts). The heterogeneity profiles (*i*.*e*. change in Hill numbers with heterogeneity order *q*) was qualitatively similar for all three measures of heterogeneity. Species presence or absence (*q* = 0) varied most strongly across groups (habitats, environments), whereas those that differentially weight common and rare taxa (*q* > 0) suggested more similarity between some groups.

For all values of *q*, network heterogeneity of node abundance almost always differed between habitats. Exceptions included comparisons between animal and plant microbiomes for *q* =1 (*p* = 0.36) and between saline and non-saline surface waters for *q* = 0 (Tables S9, S10). Similarly, network heterogeneity of link weights and overall heterogeneity differed among nearly all habitats for *q* = 0, but for only two-thirds of them with higher values of *q*. Non-significant comparisons for link-weight heterogeneity included those between sites within animals, surface waters, and sediments (Table S9). Non-significant comparisons for overall network heterogeneity were most pronounced between host-associated and free-living systems, and between saline and non-saline ones (Table S10).

#### Scaling network heterogeneity with Taylor’s Power Law

We used Taylor’s Power Law of Networks (TPLoN; Ma 2025) to examine the scaling of weighted species connectedness (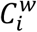; Eqn. 3) across network nodes in the EMP dataset. Although we fit the mean (*M*) and variance (*V*) of 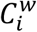 using the classic Taylor’s power law model (Eqn. 9), the scaling parameters (*b, a*, and *M*_0_) using TPLoN reflect heterogeneity-scaling relationships in complex network systems. We found significant power-law scaling across all groups and group × site combinations in the EMP; slopes (*b*) ranged from 1.462 in host-associated habitats to 2.003 in the animal corpus (Fig. 2, Table S11).

**Figure 2.**
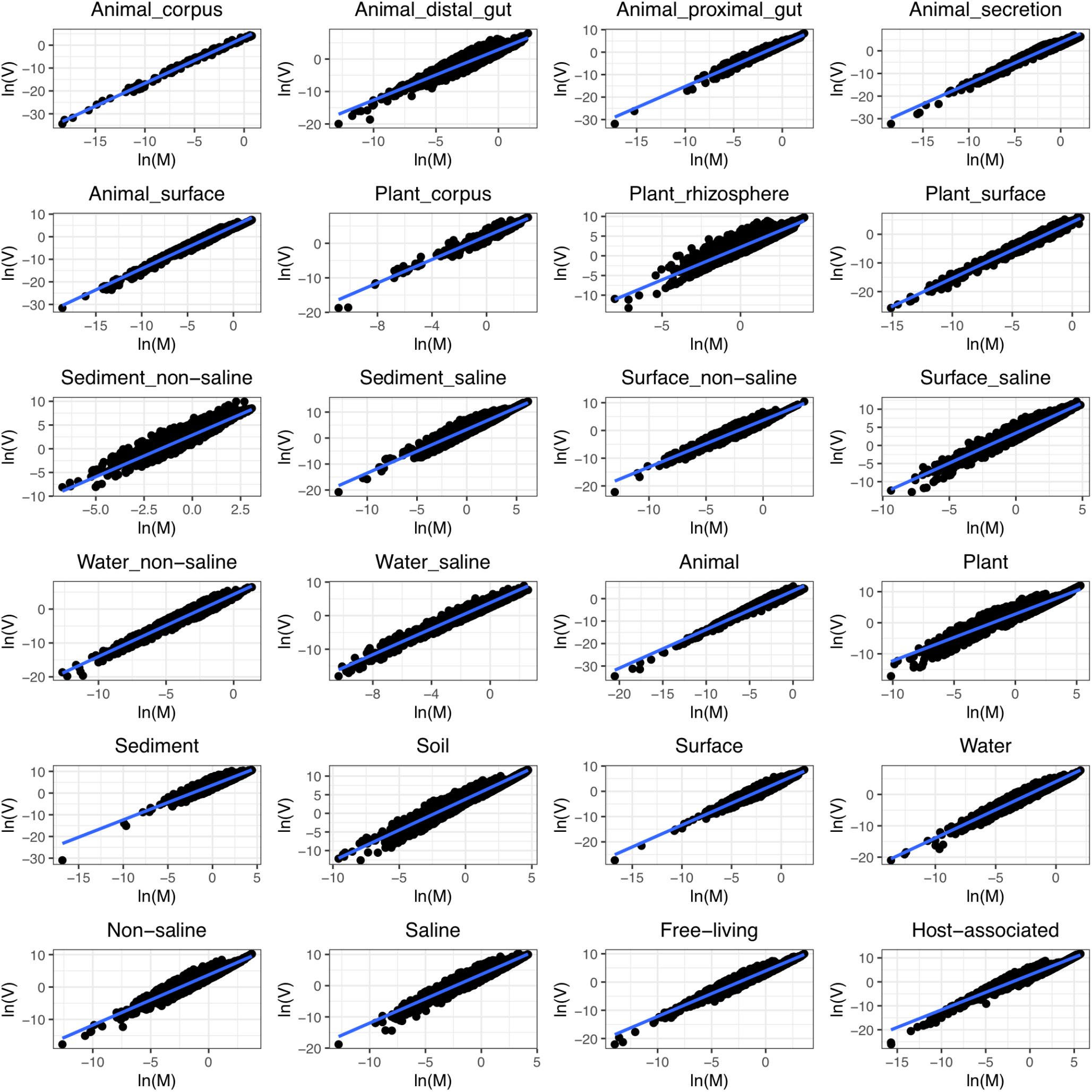
Taylor’s Power Law of Networks (TPLoN) fitted to the networks of 10 groups and 14 group × site combinations in the Earth Microbiome Project (EMP) dataset.

Among groups, microbial networks associated with plants had the lowest values of *b* (1.506), animal-associated networks were intermediate (*b* = 1.719), and free-living microbes in aquatic groups were the highest (*b* = 1.801) (Table S11). Results of permutation tests of the TPLoN parameters and *M*_*0*_ were consistent among comparisons of groups and group × site combinations (Tables S12, S13).

#### Compositional changes between networks

The preceding analyses examined heterogeneity of nodes, link weights, and entire networks. Although these aggregated measures provide information about heterogeneity at the network level, we can also look at differences in heterogeneity properties of individual OTUs that occur in multiple networks to detect “important” species (e.g. dominant, foundation, keystone species) and functional guilds.

To compare the heterogeneity of an individual species or OTU between two networks, we first determined its node heterogeneity in each of two networks *i* and *j*. We then created a 3 × 3 table (3 possibilities of heterogeneous, random or regular node in each network) of possible node “switches” for comparing networks *i* and *j* (Table 2). Among the nine possible node switches, six are *categorical* changes in heterogeneity and three are not. To analyze the node switches, we used bootstrap validation (Appendix A) to determine the heterogeneity type (category) of each node followed by permutation tests (Appendix D) to identify OTUs whose heterogeneity significantly increased or decreased from one network to another (an example is shown in Fig. 3).

**Table 2.**
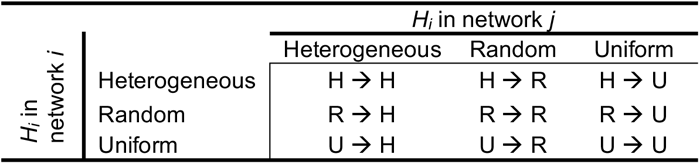
Possible switches in heterogeneity category (type) of a given node *H*_*i*_ between two networks. See Appendix D for additional details on permutation tests for node switches.

**Figure 3.**
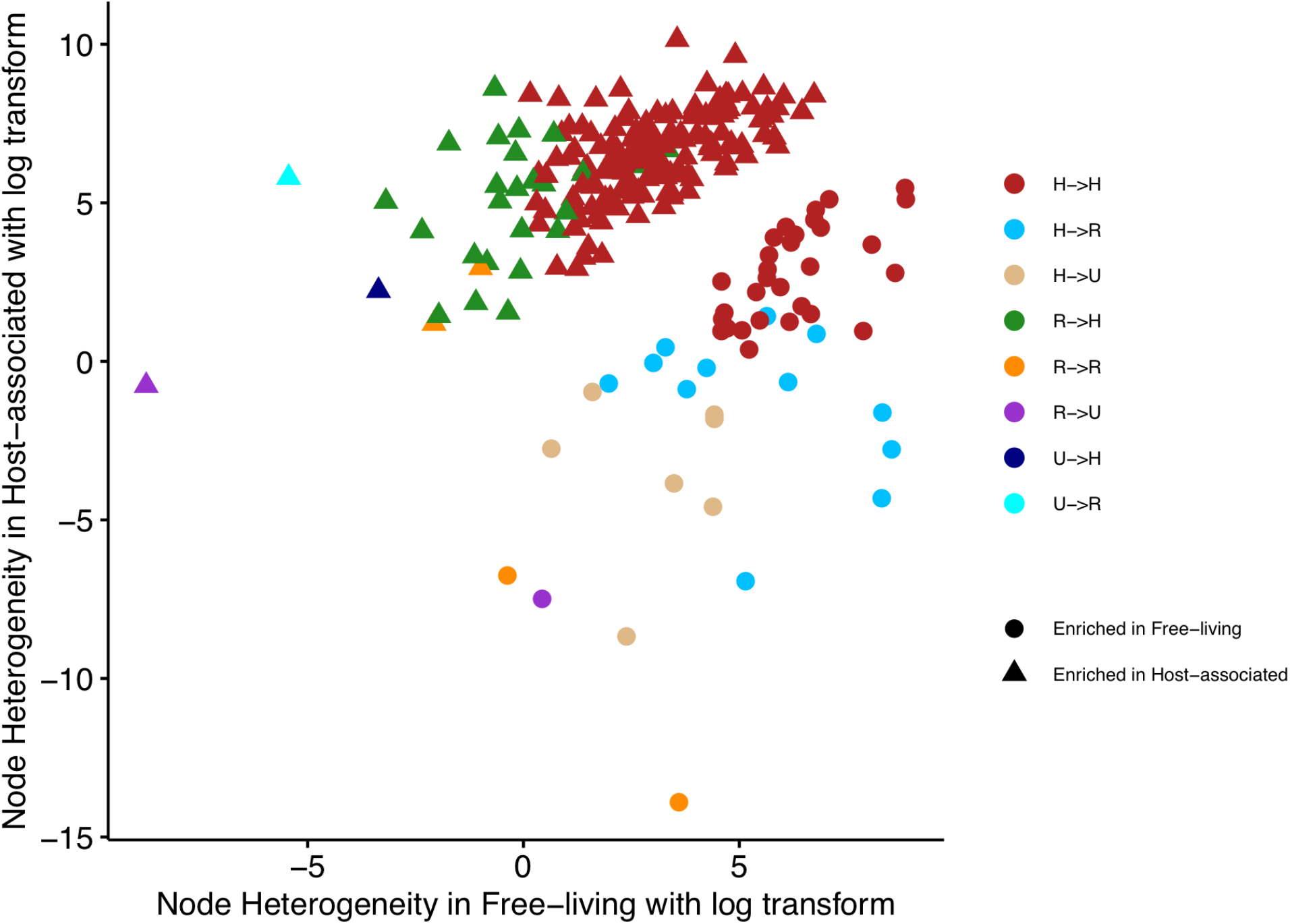
Scatter plot of OTUs (nodes) showing changes differences in node-heterogeneity of free-living and host-associated microbiome networks. Circles represent nodes with higher values of heterogeneity in free-living microbiomes; triangles indicate nodes with higher values of heterogeneity in host-associated microbiomes. Node colors denote different node switches (note that the U→U switch does not occur in this dataset). Note the clear separation of nodes enriched in each network along the diagonal.

Table 3 summaries the 4,196 node switches observed in all pairwise comparisons of networks of the groups and group × site combinations in the EMP dataset (Table S1). Only 30% of these involved switches from one heterogeneity type (heterogeneous, random, uniform) to another (off-diagonals in Table 2). Of these, 39% were H → R, 30% were R → H, 12% were H → U, 14% were U → H, and both R → U and U → R accounted for 3% each. Finer-grained details of these results are given in Tables S14 and S15; raw data used to construct these tables are in Datasets 3 and 4 (with associated metadata in Tables S16 and S17).

**Table 3.** Numbers of significant node heterogeneity switches (as in Table 2) in pairwise network comparisons of all 10 groups and 14 group × site combinations in the EMP dataset (Table S1). Switches here include actual node-switches in heterogeneity type (off-diagonals in Table 2) and nodes that remain in the same state (diagonals in Table 2). Values are summarized from finer-grained data given in Tables S14 and S15, and raw data in Datasets 3 and 4 (metadata in Tables S16 and S17).

| Switch | Number of significant alternations in heterogeneity-increased species in Network #1 | Number of significant alternations in heterogeneity-increased species in Network #2 |
| --- | --- | --- |
| **Node switches: off-diagonals in Table 2** |  |  |
| $H \rightarrow R$ | 482 | 6 |
| $H \rightarrow U$ | 148 | 0 |
| $R \rightarrow H$ | 1 | 372 |
| $R \rightarrow U$ | 32 | 4 |
| $U \rightarrow H$ | 0 | 170 |
| $U \rightarrow R$ | 0 | 44 |
| *Total node switches* | 663 | 596 |
| **Node non-switches: diagonals in Table 2** |  |  |
| $H \rightarrow H$ | 1213 | 1686 |
| $R \rightarrow R$ | 14 | 13 |
| $U \rightarrow U$ | 3 | 8 |
| *Total node non-switches* | 1230 | 1707 |

## DISCUSSION

The key distinction between diversity and heterogeneity is that the former merely enumerates distinct types (with counts optionally weighted by relative abundance) within a population, whereas the latter is a property of a collective—a group characterized by joint processes and internal structure—that quantifies the relationships among its members (Shavit & Ellison, 2021; Zelnik et al., 2023). Our methods for estimating, testing, and inference about network heterogeneity successfully identifies and quantifies these relationships. These methods are applicable to parts of networks (nodes, edges, subnetworks), scalable to entire networks and collections of networks, and are not dependent on the type of network being analyzed (e.g. microbiomes, bipartite networks, multitrophic food webs, connected patches in a landscape).

Individual elements of our method are drawn from, or inspired by, many existing approaches. For example, our measure of node heterogeneity, *H*_*i*_ (Eqn. 4), includes the node degree, *D*_*i*_, which is used in computing the degree difference between two nodes and contributes to estimates of the structural heterogeneity of complex networks (Farzam et al., 2020). Our first approach to estimating link-weighted sums (Eqn. 2) was equivalent to the measure of node strength that Bertolero et al. (2017) used to identify rich clubs. Like these (and other) approaches, our methods for estimation and inference take advantage of the information content of heterogeneity rather than considering it to be simply a reflection of heteroscedasticity or other noise in the data (Kolasa & Rollo, 1991; Sparrow, 1999).

### Scaling heterogeneities

Applying our methods to nested networks in the EMP dataset (i.e. comparisons between group × site combinations and between groups) revealed that scaling of heterogeneities (λ in Eqn. 7) in larger networks (e.g. EMP groups such as “Animals”) was different from the scaling of their subnetworks (e.g. “animal corpus”, “animal distal gut”). Heterogeneity among larger networks likely reflects differences between habitats (e.g. animals vs. plant microbiomes, free-living assemblages in sediments vs. open water), whereas heterogeneity among subnetworks within a large network reveals heterogeneity within habitats (e.g. different microbiomes within organs of animals). Creating larger groups by pooling data from subnetworks also amplifies measured heterogeneity by integrating disjoint structures, interactions among OTUs, and abundance distributions across subnetworks, thereby increasing overall variability. Finally, structures and properties of large networks also tend to reflect those of its dominant or heavily sampled subnetworks rather than the variation among all the subnetworks or poorly sampled ones. For example, the scaling of heterogeneity with animals (λ = −0.22) is the same as that of the large animal corpus microbiome, but distinct from the smaller distal gut microbiome (λ = −0.38). In the EMP dataset, these differences are most clearly apparent in host-associated microbiomes vs. free-living microbial assemblages (Fig. 3; Table S15). Understanding these differences help clarify why ecological scaling relationships are observed and increase interpretability of measures of heterogeneity at different levels of biological organization. Our results suggest that hosts impose global constraints through physiological specialization, whereas free-living conditions promote ecological opportunity—and hence higher heterogeneity—because of niche partitioning.

### Strengths and weaknesses of different measures of heterogeneity

The *V*/*M* ratio has been used extensively to properties grouped with the general term “heterogeneity” (Ma & Ellison, 2026). For example, *V*/*M* quantifies spatial aggregation in ecology, dispersion in statistics and the “Fano factor” in physics and signal processing (Harrison et al. 2008, Taylor 2019). Our analysis of the EMP dataset supports the utility of *V*/*M* for measuring and distinguishing node-level heterogeneities. However, *V*/*M* is less effective for discriminating between entire networks (Tables S8–S10), where it systematically fails to detect ecologically meaningful differences in heterogeneity among groups. In contrast, Hill-number measures, which are based on entropies rather than statistical moments, provide more interpretable measures of observed heterogeneities.

### Directions for future work

#### What is the relationship between heterogeneity and diversity?

Empirical evidence consistently identifies environmental heterogeneity (as spatial variation in habitats) as a primary driver of biodiversity (e.g. Stein et al., 2014; Udy et al., 2021). However, fragmentation that increases spatial heterogeneity beyond a certain point can reduce species diversity, resulting in a unimodal heterogeneity-diversity relationship rather than a strictly positive relationship (Allouche et al., 2012). Eishenhauer et al. (2023) suggested that heterogeneity can be used to manage for biodiversity, which in turn is crucial for sustaining ecosystem functioning and stability.

In the method we propose, Hill numbers measure both diversity and heterogeneity (Table 1). When Hill numbers are used to measure node abundances across a network, aggregating them yields a measure of network diversity. But we also use Hill numbers to node-level interactions (edges); aggregating them gives a measure of overall network heterogeneity. Thus, it should be possible to investigate Eisenhauer et al.’s (2023) integrated heterogeneity–diversity–system performance (HDP) nexus (Eisenhauer et al., 2023) using the measures and methods we have developed here.

However, such investigations have at least two difficulties. First, network heterogeneity is a measure of the entire system (or a metacommunity), so the corresponding measure of diversity should be gamma, not alpha diversity. Yet, alpha diversity is measured more frequently and gamma diversity is derived from its relationship—ironically, through “beta diversity”, which is often considered to be another kind of ecological heterogeneity; Ellison, 2010; Ma & Ellison, 2026). Care must be taken to avoid duplicating measures of heterogeneity in both sides of the analysis. Second, the parameters used to quantify heterogeneity and diversity must be of comparable types and units to permit meaningful comparisons; for example, a measure of heterogeneity or diversity and a heterogeneity scaling parameter are fundamentally different kinds of measures.

Different proxies of heterogeneity may provide an initial work-around to allow for reasonable analyses of the relationship between heterogeneity and diversity. For example, if a community-wide measure of weighted species interactions is used as a proxy for (network) heterogeneity, then the heterogeneity-diversity relationship could be tested with it. Comparison of scaling coefficients using Taylor’s power law (L. R. Taylor, 1961, 1984; R. A. J. Taylor, 2019) and its extensions (e.g. Ma, 2015, 2025; Ma & Taylor, 2020) occupy a particularly significant position. As a near-universal scaling law between variance and mean (Taylor 2019; Ma & Taylor 2025), TPL could provide a powerful and generalizable method for quantifying heterogeneity and relating it to diversity.

#### Testing the heterogeneity-stability relationship (HSR)

Heterogeneity is widely assumed to confer stability in ecological systems (e.g. Holt & Hassell, 1993; but see Sora et al., 2025). The EMP lacks the longitudinal time-series data necessary for direct tests of HSRs, but we suggest that linking our measures of heterogeneity with stiffness, which measures the resistance of a system to deformation when subject to external forces (Chen et al., 2025), could be used to test the HSR in different networks. In particular, the stiffness matrix *K*_*nXn*_ = [*k*_*ij*_] (in which *k*_*ij*_ are elastic moduli) used to determine the distribution of forces in physical structures is mathematically equivalent to the weighted adjacency matrix *W*_*nXn*_ = [*w*_*ij*_] in network science (where the *w*_*ij*_’s are edge weights); all that is required is substituting *w*_*ij*_ for *k*_*ij*_. With this substitution, the stiffness matrix can be used to quantify network resilience: node-level “stiffness” (*S*_*i*_ = Σ_*j*_ *w*_*ij*_) measures local resistance to perturbation and *K*_*nXn*_ characterizes global network stability. Chen et al. (2025) demonstrated this linkage through studies of microbiome resilience, but did not discuss microbiome heterogeneity.

Future investigation of HSRs in networks should: ensure that all co-occurrence networks are fully weighted (abundances and interactions); systematically examine correlations between heterogeneity metrics and stiffness-based stability measures; and apply shared species analysis to identify overlapping node sets that simultaneously regulate both heterogeneity and stability. The goal here would be to determine the degree of functional overlap between heterogeneity-controlling and stability-governing nodes and whether common “driver” nodes coordinate both system properties.

We note that examining HSRs in networks also entails using temporal network analysis (Holme & Saramäki 2013). Thus, any method of constructing temporal networks from co-occurrence data needs to include time-dependent edge weights, where connection strengths between species [*w*_ij_(*t*)] vary across sampling intervals. Second, linking measures of heterogeneity and stiffness must allow for time-varying measures of *V*/*M* (e.g. as *V*_*i*_(*t*)/*M*_*i*_(*t*)) ratios and stiffness matrices [*K*_n×n_(*t*)]. Finally, correlation analyses between heterogeneity and stability must account for temporal autocorrelation. All three of these requirements pose nontrivial computational and theoretical challenges, particularly in defining appropriate null models for time-varying temporal networks and establishing significance thresholds for dynamic correlations. Such work should build on established techniques for dynamic community detection (e.g. Pilosof et al., 2017) and multilayer network stability analysis (Gilarranz et al., 2017).

## CONCLUSION

Our analysis of the EMP data using our new method for estimating, testing, and making inferences about network heterogeneity support established principles of host filtering, species sorting, and environmental niche partitioning among microbiomes and free-living microbial networks. These results suggest that our measures of node, edge, and overall network heterogeneity can be used to better understand fundamental tradeoffs between functional specialization and heterogeneity in ecological networks, and illuminate how evolutionary and ecological factors jointly shape both topology and heterogeneity in ecological communities. Finally, the scaling relationships using Hill numbers and extensions of Taylor’s power law could be developed further as a quantitative tool for better understanding relationships between heterogeneity, diversity, and stability.

## Supporting information

Online Supplementary Info (CSV files)

## SUPPLEMENTAL MATERIAL

## APPENDIX A. Computational algorithm for bootstrap tests for node heterogeneity

We use a bootstrap test to filter artifacts and estimate node heterogeneity, and to partition nodes into heterogeneous, random, and uniform clusters. Each node undergoes two bootstrap tests, one for *H*_i_ > 1 and another for *H*_*i*_ < 1. Nodes failing both tests are classified as random. The computational algorithm for doing these bootstrap tests is

1. Use the WGCNA package in R (Langfelder & Horvath 2008) to calculate the weighted network from an *m* × *n* “site × species” matrix (or sample × OTU matrix as in our EMP case-study) whose entries are species abundances.
2. Compute *H*_*i*_ (heterogeneity for each node *i*) from Eqn. 4 in main text 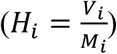, where *V*_*i*_ and *M*_*i*_ are the variance and mean of the vector of weighted species connectedness of species *i* given in Eqn. 3 in main text: 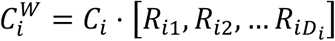. In this equation, *C*_*i*_ = *D*_*i*_/*A*_*i*_ is the connectedness of an individual species *i* in a network (*D*_*i*_ is the node-degree of species *i* and *A*_*i*_ is the mean abundance of species *i* across all the samples used to construct the network [Ma, 2025]), and [*R*_*ij*_] is the vector of interaction strengths between species *i* and *j* ∈{1, 2, …, *D*_*i*_}. Note that 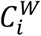 is a scalar product (and hence, a vector).
3. Resample the site × species matrix *with replacement* (at least 1000 times).
4. Generate weighted networks and compute *H*_*i*_ for each of the bootstrapped samples.
5. Compute the empirical *P* value of *H*_*i*_ for each node *i*; note that there are distinct *P* values for whether *H*_*i*_ is > 1 (heterogeneous) and < 1 (uniform): 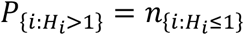 and < 1 (uniform): 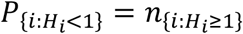. Nodes that are neither heterogeneous or uniform are “random”.

## APPENDIX B Estimating the power-law parameters for node heterogeneity

We use maximum likelihood estimation (MLE) with Kolmogorov-Smirnov (KS) optimization (Clauset et al., 2009) to estimate the scaling exponent α and lower bound or threshold *x*_*min*_ for the power-law distribution for node-level heterogeneity values (*V*_*i*_*/M*_*i*_) across the network (Eqn. 8 in the main text, here re-written as Eqn. A1):

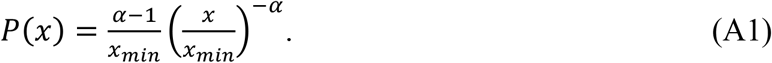

Following Clauset et al. (2009), the fitting procedure is:

1. **Estimate** *x*_*min*_: Choose the value that minimizes the KS statistic between the empirical data and power-law model for x ≥ *x*_*min*_.
2. **Calculate** α using MLE

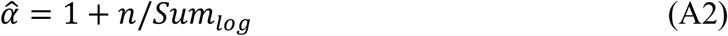

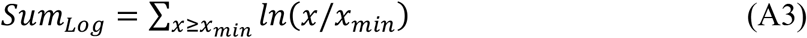

where *n* is the number of observed values satisfying x ≥*x*_*min*_.
3. **Goodness-of-Fit Test**: Generate synthetic power-law datasets with 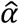 and *x*_*min*_. Compute the KS statistic for each synthetic dataset.
4. **Calculate *P* value** as the fraction of synthetic KS distances that exceed the empirical KS distance. The power-law distribution is a plausible fit if *P* ≥0.1.

## APPENDIX C Comparing heterogeneities between networks

We are interested here in comparing the scaling exponents (α) of the power-law distributions fit to more than one network and to holistically assess differences in curvature, truncation, or mid-range behavior between networks in the entire power-law distribution that may reflect network specific or distinctive patterns in heterogeneity.

We use a *Z-*test to compare scaling exponents (α_1_, α_2_) of two power-law distributions:

1. Estimate α_1_, α_2_, and their standard errors (σ_1_, σ_2_) using MLE;
2. Compute the Z-test statistic

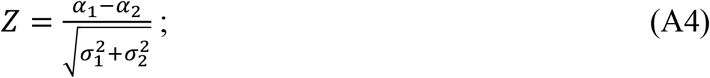
3. Reject the null hypothesis (*α*_1_ = *α*_2_) if |*Z*| > 1.96 (*P* < 0.05).

The limitation of the Z-test is that it only tests the scaling parameter α, but ignores other distributional differences. Using a Z-test for comparing the other parameter of the power-law distribution, *x*_*min*_, is not recommended because *x*_min_ is typically estimated with K-S minimization (Appendix B, above), which does not have an analytically-determined standard error. However, for completeness, we indicate how a Z-test could be used to compare two values of *x*_min_ using a bootstrap to estimate their standard deviations:

1. Resample each dataset with replacement (usually 1000 bootstrap samples);
2. Estimate *x*_min_ for each bootstrap sample using K-S minimization;
3. Compare the 95% confidence intervals (CIs) of the *x*_min_ values for each distribution. Reject the null hypothesis that they are equal if the two CIs do not overlap.

A Kolmogorov-Smirnov test can be used to compare the shapes (not just the scaling parameter) of two empirical distributions:

1. Compute the cumulative distribution functions (CDFs) for both samples;
2. Calculate the maximal vertical distance (*D*) between CDFs:

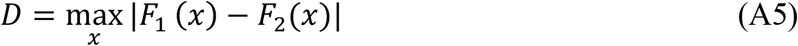
3. Reject the null hypothesis (the two distributions are identical) if *D* exceeds its critical value, which depends on sample size (see Chapter 2 of Dormann & Ellison, 2025)

The Kolmogorov-Smirnov (K-S) test detects differences in the complete shape of empirical distributions, including variations in curvature, truncation points, or other structural features beyond just the scaling exponent. In contrast to the Z-test—which is designed specifically to compare power-law scaling parameters (α)—the K-S test offers three key advantages: First, it is non-parametric, meaning it does not assume the data follows a power-law or any other specific distribution. Second, it evaluates dissimilarities between distributions holistically, capturing deviations across their entire range rather than focusing solely on tail behavior. Third, although the K-S test is less sensitive than the Z-test to tail-specific differences in α, it excels at identifying broader distributional disparities that may arise from noise, thresholds, or mixed generative processes.

## APPENDIX D Permutation tests to compare heterogeneities between networks

This permutation algorithm serves two key purposes: (1) Identifying nodes with significant changes in heterogeneity, including nodes that switch between clusters and those that change their heterogeneity significantly but remain within a particular cluster; and (2) detecting global differences in heterogeneity between two networks. The permutation test is implemented with the following general algorithm:

1. For a pair of networks represented by an *m × n* (site × species or sample × OTU matrix), construct a weighted network for each treatment using the WGCNA package in R (Langfelder & Horvath 2008).
2. Compute their node heterogeneity (*H*_*i*_), network heterogeneity (^*q*^*H*), TPLoN network heterogeneity scaling parameters (*a, b* from *V = aM*^*b*^) (full equations in Table 1 of main text), and the heterogeneity criticality threshold (HCT): 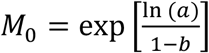 (Eqn. 11 in main text).
3. Compute the absolute difference (Δ) between the two networks for each measure of heterogeneity or TPLoN parameter, where for measure or parameter *η*, Δ_*η*_ = |*η*_1_ - *η*_2_|.
4. Pool together all samples from the two networks, and randomly permutate the orders (indices) of all samples, *i*.*e*., generating the total permutations of all samples from both networks. For example, if there are *r* samples in in the dataset used to building one network and *s* samples in the dataset used to build the other network, the number of possible total permutation is (*r* + *s*)!.
5. Randomly select 1000 orders out of the total permutations [(*m*+*n*)!] from step 4. For each of the selected orders, designate the first *r* samples as belonging to dataset 1 for network 1 and the remaining *s* samples as belonging to dataset 2 for network 2.
6. Repeat steps 1 – 3 for each of the selected orders, generating 1000 pairs of weighted networks and corresponding 1000 pairs of heterogeneity measures and parameters η, respectively: Δ_*pη*_*=*|*η*_*p1*_*–η*_*p2*_|, where the subscript *p* notes ‘*permutation*’. A total of 1000 _*pη*_ is obtained in this step. The computed parameter differences (Δ_*pη*_: Step 3) include differences in heterogeneity of both individual nodes and entire networks.
7. Compute the means and standard deviations of the model parameters from the 1000 permutations, as well as the empirical *P* value for the permutation test. This *P* value is defined as the number of permutations with Δ _*pη*_ > Δ_*η*_, divided by the total number of permutations (1000). If *P* ≤ 0.05, we infer that the measurement or parameter η differs significantly between the two networks.

For testing differences in node heterogeneity, we compute the absolute difference in node heterogeneity between two networks (Δ = |*H*_1_ − *H*_2_|) and its associated *P* value, using the same algorithm applied to network-level parameters. The decision rules for node heterogeneity are based on two observable changes in pairwise network comparisons:

1. *Directional change in heterogeneity*—For each species, if *h*_1_ > *h*_2_, the node’s heterogeneity significantly increases in network 1 and decreases in network 2 (or *vice versa*). In other words, each comparison yields three classifications: *raised nodes* (heterogeneity increased in network 1, equivalent to decreased nodes in network 2), *declined nodes* (heterogeneity decreased in network 1 and increased in network 2), and *stable nodes* (no significant change in node heterogeneity in the two networks relative to each other).
2. *Cluster membership changes*: Nodes that switch clusters (e.g., from *h*_1_ > 1 to *h*_2_ = 1) are classified as transformed. Recall from Appendix A, above, that the bootstrap test identified nodes as being either heterogeneous, uniform, or random. Thus, the permutation test can identify six possible transformations (between heterogeneous [H], uniform [U], and random [R] states) in pairwise comparisons (H→R, H→U, R→H, R→U, U→R, U→H), along with three cases (H→H, R→R, U→U) in which node type is not altered.

To simplify network comparison, we excluded nodes showing no significant heterogeneity changes between networks and omitted OTUs absent in either network being compared because of treatment-specific absence, removal by network construction algorithms, or elimination as spurious reads (e.g., singletons). Although comparisons involving the omitted OTUs could yield biological insights, their analysis presents technical complications. For example, when an OTU is absent in one network, its node heterogeneity = 0, making formal algorithmic application trivial: the analysis would merely test whether the non-zero heterogeneity in the counterpart network is statistically significant.

In summary, the permutation test algorithm systematically categorizes node-level heterogeneity differences while accounting for directional changes, cluster transitions (membership switches), and final classification statuses.

## Acknowledgments

This work was supported by a Charles Bullard Fellowship in Forest Research to ZM.

---

**Table S1.** Summary characteristics of the 23,727 samples from the Earth Microbiome Project (EMP) dataset (after Thompson et al., 2017, Nature).

| Group (environment, habitat, or host) | Site | Number of Samples | Notes |
| --- | --- | --- | --- |
| Animal ($N=9071$) | Corpus | 328 | Samples of bees, flies, includes whole head, whole larva, whole pupa, whole gut and royal jelly. Effect of Tetracycline application. |
|  | Distal gut | 4158 | Primarily feces samples from Human, Dog, Cat, Fox, Swamp Wallaby, Emu, Green Iguanas, Myrmecophagous Mammals (anteaters, armadillos, etc.), Cows, Cape Buffalo, Phyllostomid Bats, Hibernating Bears, Spider Monkeys, Seba's Short-Tailed Bats, Colobine Primates. |
|  | Proximal gut | 367 | Mucosal samples from stomach compartments of colobine primates; Gizzard and upper intestine samples from birds. |
|  | Secretion | 1257 | oral/mouth samples; saliva |
|  | Surface | 2961 | Primarily skin samples from Human skin (palms), Fish mucus, Bird eggshells, Humpback Whale skin, Fish, Frog, Rat, Reptile skin. |
| Plant ($N=2287$) | Corpus | 125 | Internal plant tissues or specific structures, Plant corpus (unspecified), Flowers, grapes, leaves. |
|  | Rhizosphere | 554 | Soil closely associated with plant roots. |
|  | Surface | 1608 | Biofilms on plant surfaces |
| Sediment ($N=1103$) | Non-saline | 544 | Freshwater sediments. Specific contexts include: River sediment, Postglacial ponds, Tibetan Plateau lakes |
|  | Saline | 559 | Marine sediments. Specific contexts include: Coastal marine (sandy-mud), Carbonate chimneys/seeps, Methane hydrate-bearing deep marine, Mission Bay, Deepwater Horizon spill site, Polluted polar coastal, Tibetan Plateau alkaline lakes. |
| Soil ($N=4279$) | Non-saline | 4279 | Terrestrial soil samples. Specific contexts include: Corn experimental field, Semi-arid steppe, Hydrocarbon-contaminated Antarctic, Upland Alaskan boreal forest, Volcanic, Post-glacial pond profiles, Urban Manhattan, Alaska peat, Changbai Mountain, African farms, Malaysian forest/plantation, Nicaragua coffee, Panama precipitation gradient, Biochar-amended agricultural, Recovering crusts in Glen Canyon |
| Surface ($N=1388$) | Non-saline | 1271 | Non-saline surfaces (non-animal, non-plant). Includes: Komodo Dragon habitat surfaces, Indoor surfaces (e.g., house floors). |
|  | Saline | 117 | Saline surfaces (non-animal, non-plant). Includes: Rocky shores, Carbonate chimneys (Lost City Hydrothermal Field) |
| Aquatic ($N=5599$) | Non-saline | 4915 | Freshwater samples. Specific contexts include: Lakes (e.g., Pavilion Lake, German lakes, Lake Mendota, North temperate bog lake), Rivers (e.g., Colorado River), Cenotes (Yucatan) |
|  | Saline | 684 | Marine/Seawater samples. Specific contexts include: Coastal pelagic mesocosms, Seasonal marine dynamics, Catlin Arctic Survey, Mediterranean Sea, Cenotes (Yucatan), Deep North Atlantic. |
| Free living communities ($N=12,369$) | Non-saline | 11009 | Combined all samples of non-saline communities from sediment, soil, surface, and aquatic groups |
|  | Saline | 1360 | Combined all samples of saline communities from sediment, soil, surface, and aquatic groups |
| Host-associated communities |  | 11358 | Combined all samples of animal and plant communities. |

**Table S2.**
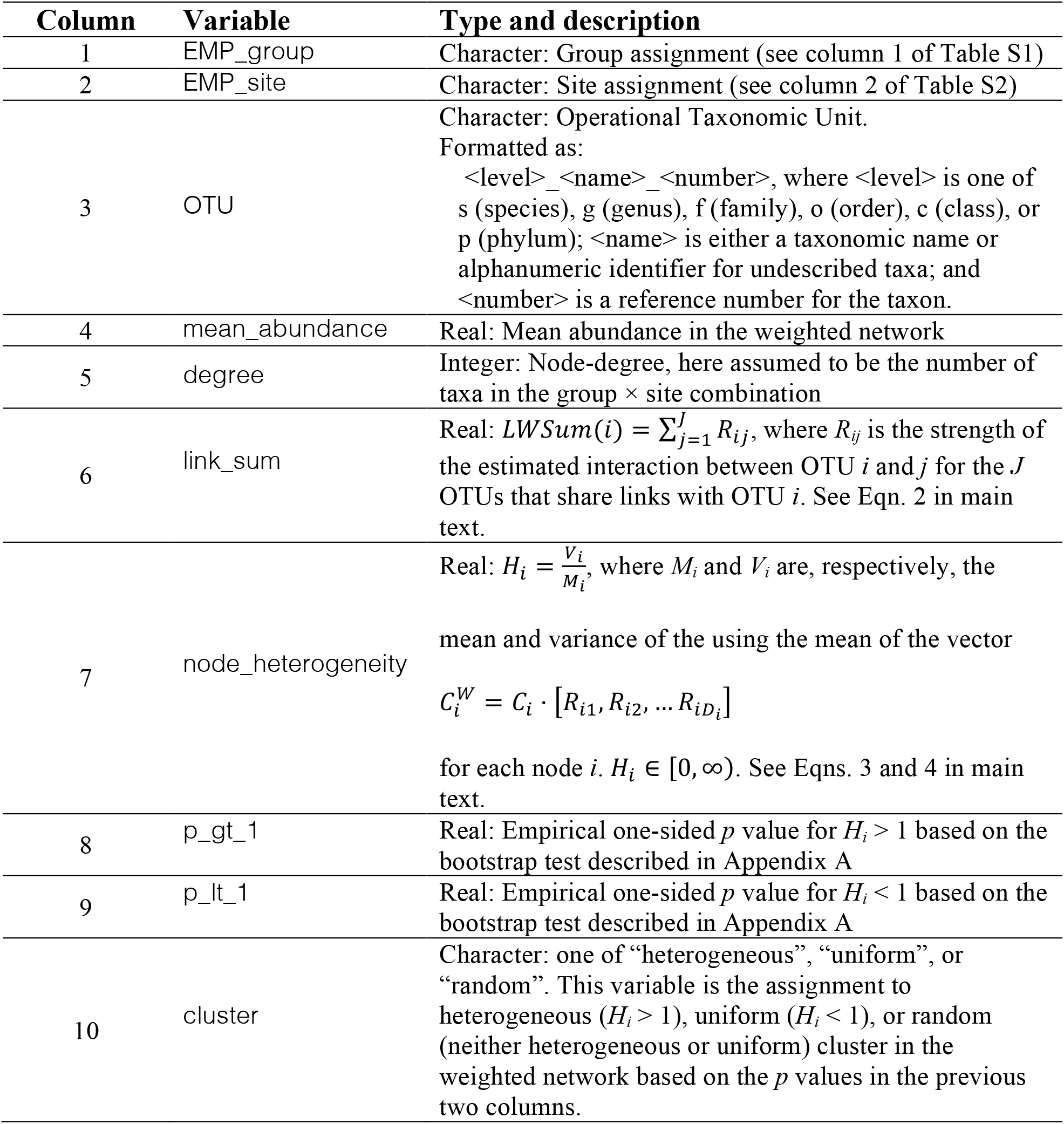
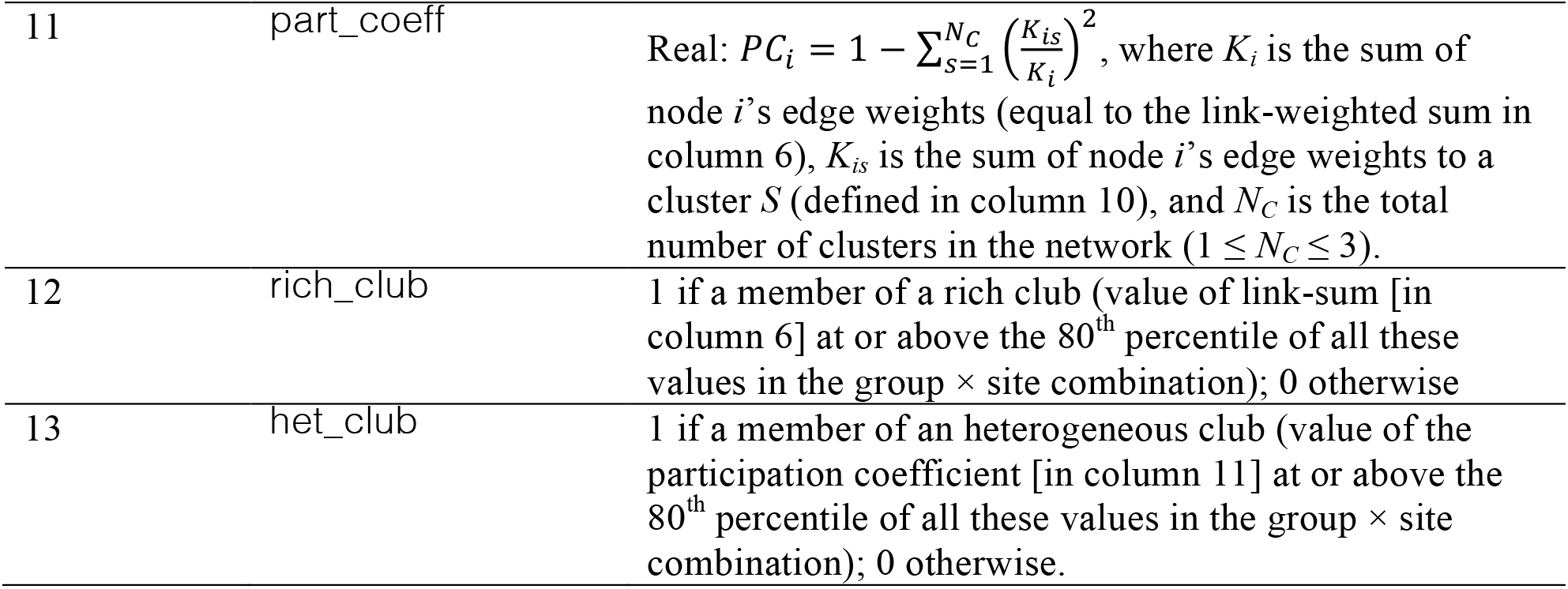
Metadata for Dataset S1. **Filename**: DatasetS1.csv **Format**: comma-separated values (.csv) **Description**: OTUs, their mean abundance, node heterogeneities, assignment of each OTU to a heterogeneity cluster (determined by p-value), link-sum (used to determine rich-club membership), participation coefficient (used to determine heterogeneous-club membership) for each of 14 unique combinations of group × site in the EMP dataset (Table S1). **Variable definitions**:

| Column | Variable | Type and description |
| --- | --- | --- |
| 1 | EMP_group | Character: Group assignment (see column 1 of Table S1) |
| 2 | EMP_site | Character: Site assignment (see column 2 of Table S2) |
| 3 | OTU | Character: Operational Taxonomic Unit.<br>Formatted as:<br><level>_<name>_<number>, where <level> is one of s (species), g (genus), f (family), o (order), c (class), or p (phylum); <name> is either a taxonomic name or alphanumeric identifier for undescribed taxa; and <number> is a reference number for the taxon. |
| 4 | mean_abundance | Real: Mean abundance in the weighted network |
| 5 | degree | Integer: Node-degree, here assumed to be the number of taxa in the group × site combination |
| 6 | link_sum | Real: $LWSum(i) = \sum_{j=1}^J R_{ij}$, where $R_{ij}$ is the strength of the estimated interaction between OTU $i$ and $j$ for the $J$ OTUs that share links with OTU $i$. See Eqn. 2 in main text. |
| 7 | node_heterogeneity | Real: $H_i = \frac{V_i}{M_i}$, where $M_i$ and $V_i$ are, respectively, the mean and variance of the using the mean of the vector $C_i^W = C_i \cdot [R_{i1}, R_{i2}, \dots, R_{iD_i}]$ for each node $i$. $H_i \in [0, \infty)$. See Eqns. 3 and 4 in main text. |
| 8 | p_gt_1 | Real: Empirical one-sided $p$ value for $H_i > 1$ based on the bootstrap test described in Appendix A |
| 9 | p_lt_1 | Real: Empirical one-sided $p$ value for $H_i < 1$ based on the bootstrap test described in Appendix A |
| 10 | cluster | Character: one of “heterogeneous”, “uniform”, or “random”. This variable is the assignment to heterogeneous ($H_i > 1$), uniform ($H_i < 1$), or random (neither heterogeneous or uniform) cluster in the weighted network based on the $p$ values in the previous two columns. |
| 11 | part_coeff | Real: $PC_i = 1 - \sum_{S=1}^{N_C} \left( \frac{K_{is}}{K_i} \right)^2$, where $K_i$ is the sum of node $i$'s edge weights (equal to the link-weighted sum in column 6), $K_{is}$ is the sum of node $i$'s edge weights to a cluster $S$ (defined in column 10), and $N_C$ is the total number of clusters in the network ($1 \leq N_C \leq 3$). |
| 12 | rich_club | 1 if a member of a rich club (value of link-sum [in column 6] at or above the 80<sup>th</sup> percentile of all these values in the group × site combination); 0 otherwise |
| 13 | het_club | 1 if a member of an heterogeneous club (value of the participation coefficient [in column 11] at or above the 80<sup>th</sup> percentile of all these values in the group × site combination); 0 otherwise. |

**Table S3.**
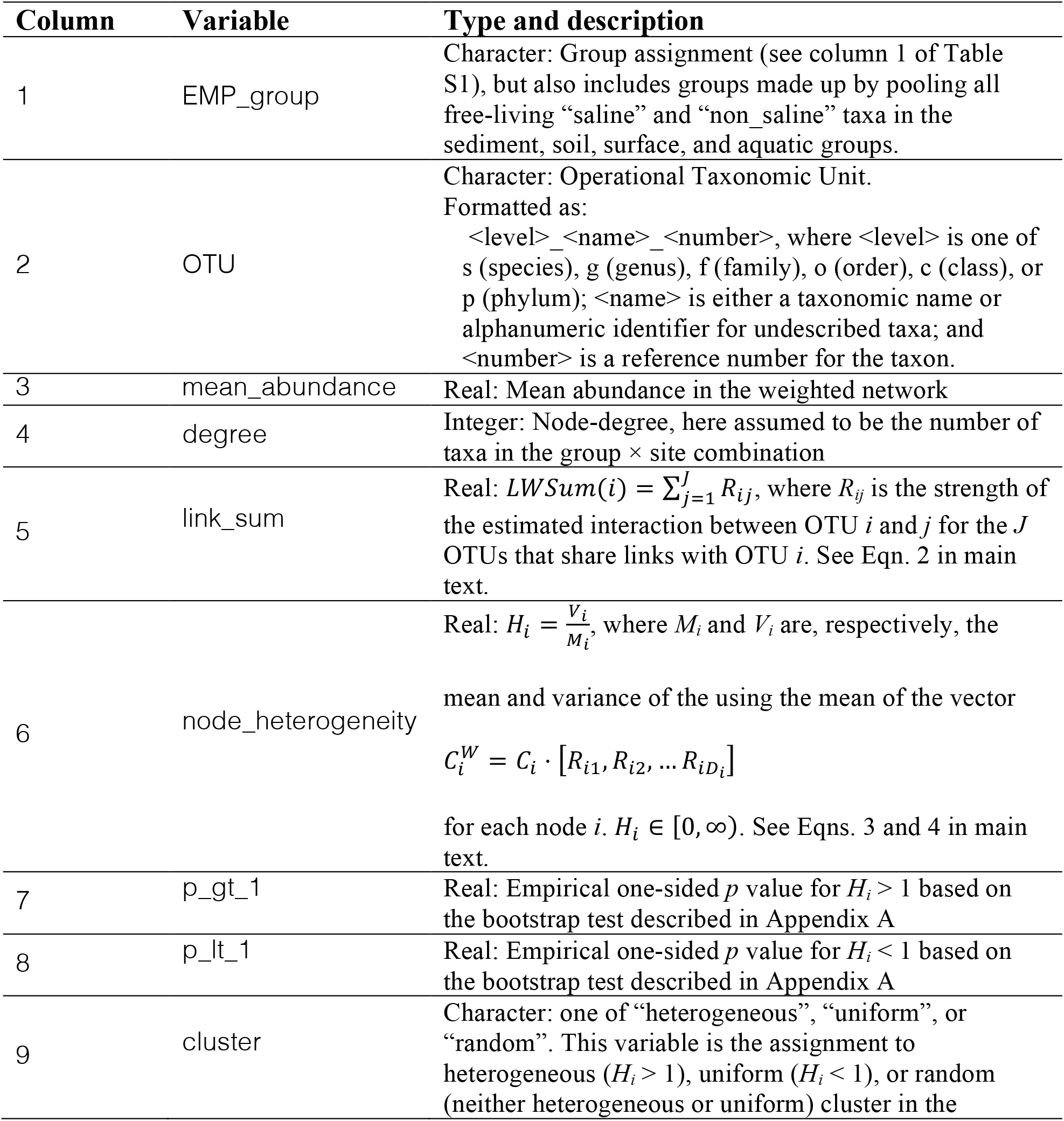
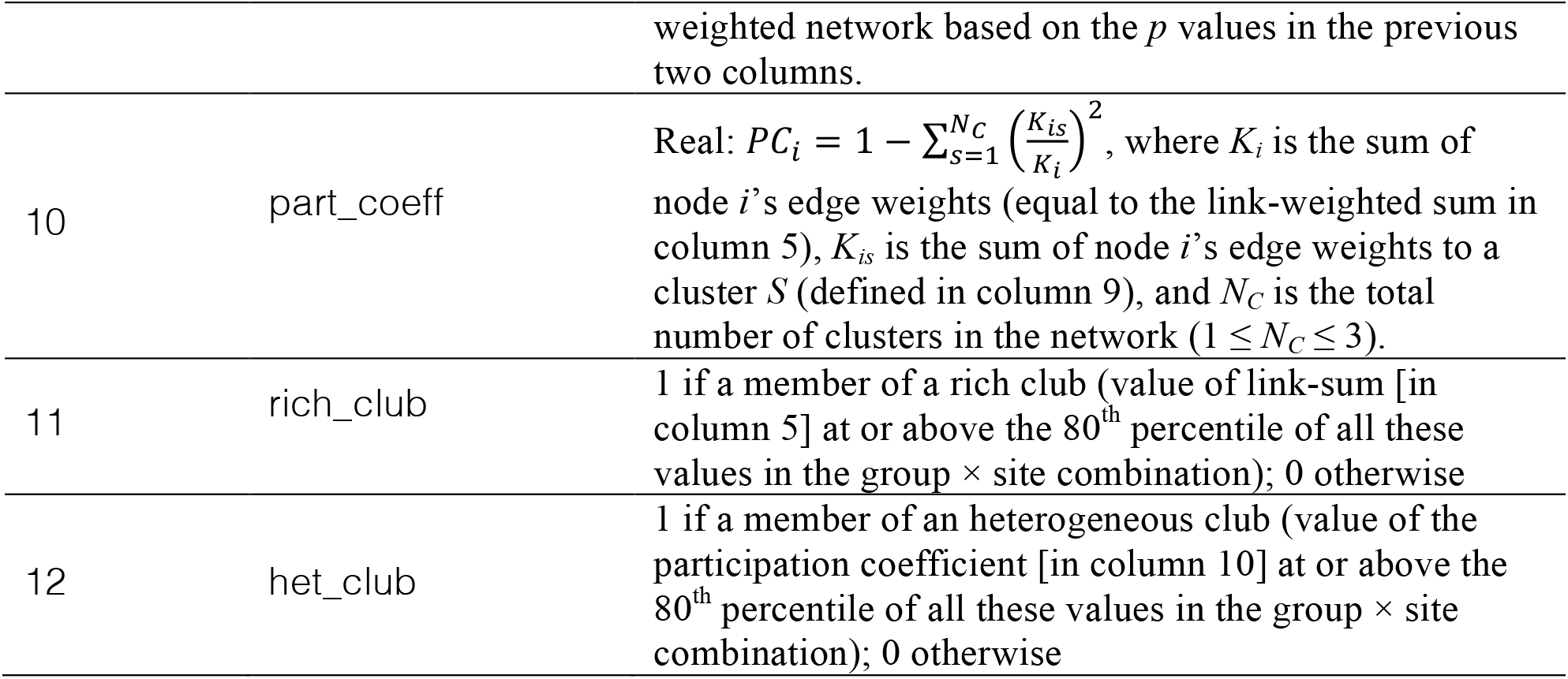
Metadata for Dataset S2. **Filename**: DatasetS2.csv **Format**: comma-separated values (.csv) **Description**: OTUs, their mean abundance, node heterogeneities, assignment of each OTU to a heterogeneity cluster (determined by p-value), link-sum (used to determine rich-club membership), participation coefficient (used to determine heterogeneous-club membership) for each of 10 unique combinations of group (pooled over sites) in the EMP dataset (Table S1). **Variable definitions**:

**Table S4.** Network characteristics of the 24 weighted networks in the EMP dataset. Values shown are the mean species (node) heterogeneity (*H*_*i*_*=V*_*i*_*/M*_*i*_), estimated percentages of heterogeneous (*P*(*H*_*i*_ > 1) < 0.05), uniform (*P*(*H*_*i*_ < 1) < 0.05), and random nodes (otherwise) for each of the group × site combination (Dataset S1) and each of the pooled groups (Dataset S2). Groups and sites as defined in Table S1. For the bootstrap-validated estimates, the mean *Hi* for each cluster (heterogeneous, uniform, and random) are also given. See Appendix A for description of the bootstrap algorithm.

|  |  | Raw data |  |  | Validated with bootstrap test |  |  |  |  |  |
| --- | --- | --- | --- | --- | --- | --- | --- | --- | --- | --- |
| Group | Site | Mean $H_i$ | Heterogeneous nodes (%) | Uniform nodes (%) | Heterogeneous nodes | | Uniform nodes | | Random nodes | |
| | | | | | % | mean $H_i$ | % | mean $H_i$ | % | mean $H_i$ |
| Animal | Corpus | 4.267 | 40.57 | 59.43 | 28.30 | 13.360 | 23.58 | 0.075 | 48.113 | 0.973 |
|  | Distal gut | 19.990 | 83.25 | 16.75 | 75.94 | 25.817 | 2.89 | 0.343 | 21.167 | 1.764 |
|  | Proximal gut | 24.569 | 84.12 | 15.88 | 77.82 | 31.315 | 6.08 | 0.217 | 16.104 | 1.166 |
|  | Secretion | 16.251 | 70.05 | 29.95 | 60.22 | 25.834 | 10.62 | 0.106 | 29.160 | 2.339 |
|  | Surface | 22.985 | 76.69 | 23.31 | 61.57 | 35.139 | 4.54 | 0.085 | 33.897 | 3.974 |
| Plant | Corpus | 18.896 | 89.22 | 10.78 | 82.04 | 22.536 | 3.29 | 0.159 | 14.671 | 2.748 |
|  | Rhizosphere | 50.337 | 97.26 | 2.74 | 95.36 | 52.701 | 0.91 | 0.217 | 3.735 | 2.132 |
|  | Surface | 12.092 | 57.80 | 42.20 | 43.69 | 24.457 | 16.96 | 0.061 | 39.349 | 3.547 |
| Sediment | Non-saline | 45.132 | 97.33 | 2.67 | 95.56 | 47.179 | 0.72 | 0.350 | 3.721 | 1.243 |
|  | Saline | 211.373 | 97.72 | 2.28 | 96.11 | 219.838 | 0.90 | 0.301 | 2.992 | 2.776 |
| Surface | Non-saline | 37.866 | 91.70 | 8.30 | 86.93 | 43.084 | 1.79 | 0.209 | 11.284 | 3.650 |
|  | Saline | 123.285 | 97.29 | 2.71 | 95.77 | 128.685 | 0.27 | 0.045 | 3.958 | 1.042 |
| Aquatic | Non-saline | 15.639 | 69.46 | 30.54 | 53.28 | 26.293 | 8.21 | 0.129 | 38.506 | 4.202 |
|  | Saline | 32.360 | 82.45 | 17.55 | 75.62 | 42.292 | 9.32 | 0.162 | 15.051 | 2.404 |
| Animal |  | 8.705 | 71.08 | 28.92 | 58.82 | 13.892 | 6.99 | 0.147 | 34.191 | 1.530 |
| Plant |  | 84.678 | 96.51 | 3.49 | 94.98 | 89.051 | 0.96 | 0.121 | 4.054 | 2.324 |
| Sediment |  | 124.777 | 99.64 | 0.36 | 99.30 | 125.648 | 0.09 | 0.374 | 0.611 | 1.736 |
| Soil |  | 84.479 | 95.86 | 4.14 | 93.14 | 90.447 | 0.54 | 0.202 | 6.317 | 3.764 |
| Surface |  | 34.703 | 92.77 | 7.23 | 89.12 | 38.658 | 1.01 | 0.228 | 9.874 | 2.525 |
| Aquatic |  | 21.877 | 81.34 | 18.66 | 71.21 | 29.887 | 5.23 | 0.217 | 23.564 | 2.477 |
| Non-saline |  | 52.165 | 97.14 | 2.86 | 95.38 | 54.415 | 0.22 | 0.298 | 4.396 | 5.938 |
| Saline |  | 83.738 | 96.70 | 3.30 | 95.01 | 88.069 | 1.06 | 0.248 | 3.926 | 1.479 |
| Free-living |  | 51.381 | 95.42 | 4.58 | 93.15 | 55.035 | 0.42 | 0.290 | 6.429 | 1.773 |
| Host-associated |  | 84.310 | 91.24 | 8.76 | 88.16 | 95.319 | 1.22 | 0.159 | 10.616 | 2.560 |
| **Mean** |  | **51.932** | **85.16** | **14.84** | **78.64** | **56.944** | **5.51** | **0.178** | **16.33** | **8.616** |
| **Standard Error** |  | **15.339** | **3.07** | **3.07** | **4.06** | **16.763** | **1.44** | **0.024** | **3.37** | **4.116** |

**Table S5.** Fit of observed distributions of node heterogeneity to power-law distributions. Values shown are estimates of parameters (α, *x*_min_) and *p* values comparing a theoretical power-law distribution with the observed distributions of node heterogeneity of the 14 group × site combinations and 10 groups of the EMP dataset (Table S1). See Appendix B for a description of the algorithm used to estimate the parameters of the power-law distributions.

| Group | Site | $\alpha$ | $x_{\min}$ | Fit ($P$-value) |
| --- | --- | --- | --- | --- |
| Animal | Corpus | 1.844 | 3.289 | 0.542 |
|  | Distal gut | 2.058 | 17.933 | 0.088 |
|  | Proximal gut | 1.936 | 13.228 | 0.001 |
|  | Secretion | 2.599 | 26.806 | 0.179 |
|  | Surface | 1.715 | 8.148 | 0.000 |
| Plant | Corpus | 3.090 | 32.638 | 0.950 |
|  | Rhizosphere | 2.279 | 41.298 | 0.000 |
|  | Surface | 1.649 | 3.145 | 0.000 |
| Sediment | Non-saline | 2.235 | 34.667 | 0.002 |
|  | Saline | 2.081 | 176.278 | 0.000 |
| Surface | Non-saline | 2.254 | 39.465 | 0.241 |
|  | Saline | 2.843 | 210.091 | 0.008 |
| Aquatic | Non-saline | 1.785 | 6.841 | 0.000 |
|  | Saline | 3.003 | 99.988 | 0.478 |
| Animal |  | 2.328 | 10.564 | 0.694 |
| Plant |  | 2.409 | 87.280 | 0.002 |
| Sediment |  | 2.189 | 88.271 | 0.000 |
| Soil |  | 1.881 | 48.299 | 0.000 |
| Surface |  | 2.453 | 46.950 | 0.115 |
| Aquatic |  | 1.858 | 10.043 | 0.000 |
| Non-saline |  | 1.969 | 26.890 | 0.000 |
| Saline |  | 2.989 | 195.463 | 0.694 |
| Free-living |  | 2.251 | 55.840 | 0.014 |
| Host-associated |  | 4.118 | 208.548 | 0.070 |

**Table S6.** Results of *Z* tests and Kolomogorov-Smirnov tests comparing power-law distributions of estimated node heterogeneities of weighted networks representing 14 group × site combinations in the EMP dataset. Values shown are *Z* and *p* values comparing node heterogeneities in Site 1 and Site 2 (estimates of α and *x*_min_ being compared are given in the first 14 rows of Table S5). See Appendix C for details of how we use the *Z* test and Kolmogorov-Smirnov test to compare power-law distributions.

| Group | Site 1 | Site 2 | $Z$ | $P(Z)$ | $P$-value from (K-S test) |
| --- | --- | --- | --- | --- | --- |
| Animal | corpus | distal gut | -11.382 | 0.000 | 0.000 |
|  |  | proximal gut | -11.804 | 0.000 | 0.000 |
|  |  | secretion | -8.751 | 0.000 | 0.000 |
|  |  | surface | -12.128 | 0.000 | 0.000 |
|  | distal gut | proximal gut | -2.479 | 0.013 | 0.000 |
|  |  | secretion | 2.449 | 0.014 | 0.000 |
|  |  | surface | -1.778 | 0.075 | 0.000 |
|  | proximal gut | secretion | 4.524 | 0.000 | 0.000 |
|  |  | surface | 0.804 | 0.422 | 0.000 |
|  | secretion | surface | -4.021 | 0.000 | 0.012 |
| Plant | corpus | rhizosphere | -17.599 | 0.000 | 0.000 |
|  |  | surface | 4.042 | 0.000 | 0.000 |
|  | rhizosphere | surface | 24.227 | 0.000 | 0.000 |
| Sediment | non-saline | saline | -28.613 | 0.000 | 0.000 |
| Surface | non-saline | saline | -23.761 | 0.000 | 0.000 |
| Aquatic | non-saline | saline | -8.562 | 0.000 | 0.000 |

**Table S7.** Results of *Z* tests and Kolomogorov-Smirnov tests comparing power-law distributions of estimated node heterogeneities of weighted networks representing 10 groups in the EMP dataset. Values shown are *Z* and *p* values comparing node heterogeneities in Site 1 and Site 2 (estimates of α and *x*_min_ being compared are given in the last 10 rows of Table S5). See Appendix C for details of how we use the *Z* test and Kolmogorov-Smirnov test to compare power-law distributions.

| Group 1 | Group 2 | Z | P-Value<br>from (Z-test) | P-value from<br>(K-S test) |
| --- | --- | --- | --- | --- |
| Animal | Plant | -35.983 | 0.000 | 0.000 |
|  | Sediment | -38.422 | 0.000 | 0.000 |
|  | Soil | -32.577 | 0.000 | 0.000 |
|  | Surface | -17.066 | 0.000 | 0.000 |
|  | Aquatic | -11.771 | 0.000 | 0.000 |
| Plant | Sediment | -11.190 | 0.000 | 0.000 |
|  | Soil | 0.066 | 0.948 | 0.000 |
|  | Surface | 20.339 | 0.000 | 0.000 |
|  | Aquatic | 28.174 | 0.000 | 0.000 |
| Sediment | Soil | 10.850 | 0.000 | 0.000 |
|  | Surface | 27.529 | 0.000 | 0.000 |
|  | Aquatic | 33.147 | 0.000 | 0.000 |
| Soil | Surface | 18.828 | 0.000 | 0.000 |
|  | Aquatic | 25.727 | 0.000 | 0.000 |
| Surface | Aquatic | 7.622 | 0.000 | 0.000 |
| Non-saline | Saline | -11.086 | 0.000 | 0.000 |
| Free-living | Host-associated | -11.330 | 0.000 | 0.000 |

**Table S8.** Measures of network heterogeneity for different Hill numbers. Network characteristics of the 24 weighted networks in the EMP dataset. Values shown are network heterogeneity for node abundances, for link weights, and overall heterogeneity for Hill numbers (Rényi entropy) *q* ∈ {0, 1, 2, 3}.

| Group | Site | Network Heterogeneity in Node Abundances |  |  |  | Network Heterogeneity in Link Weights |  |  | Overall Network Heterogeneity |  |  |  |
| --- | --- | --- | --- | --- | --- | --- | --- | --- | --- | --- | --- | --- |
| | | $q=0$ | $q=1$ | $q=2$ | $q=3$ | $q=1$ | $q=2$ | $q=3$ | $q=1$ | $q=2$ | $q=3$ | Network $V/M$ |
| Animal | Corpus | 106 | 24.989 | 16.117 | 13.344 | 47.792 | 37.996 | 33.663 | 28.685 | 20.430 | 17.485 | 18.044 |
|  | Distal gut | 1696 | 510.870 | 139.318 | 65.332 | 719.831 | 494.398 | 401.841 | 555.629 | 282.068 | 186.752 | 100.263 |
|  | Proximal gut | 888 | 223.204 | 98.853 | 66.759 | 580.505 | 438.123 | 363.022 | 358.381 | 207.526 | 142.313 | 80.652 |
|  | Secretion | 631 | 87.915 | 32.033 | 22.537 | 192.701 | 128.037 | 108.736 | 249.040 | 168.716 | 129.042 | 44.599 |
|  | Surface | 1124 | 159.123 | 46.879 | 27.655 | 289.183 | 135.615 | 102.290 | 393.444 | 248.732 | 189.689 | 80.953 |
| Plant | Corpus | 334 | 2.974 | 1.592 | 1.432 | 256.471 | 216.282 | 191.662 | 184.189 | 126.162 | 95.086 | 31.223 |
|  | Rhizosphere | 4418 | 1313.572 | 304.089 | 132.998 | 2200.563 | 1594.664 | 1347.081 | 2293.267 | 1267.978 | 743.519 | 125.081 |
|  | Surface | 737 | 101.311 | 36.956 | 24.849 | 359.259 | 265.472 | 222.230 | 202.513 | 115.673 | 81.973 | 65.038 |
| Sediment | Non-saline | 2768 | 912.415 | 374.154 | 228.282 | 1937.722 | 1481.857 | 1205.723 | 1175.829 | 493.342 | 258.442 | 208.165 |
|  | Saline | 4345 | 1129.073 | 316.265 | 148.650 | 1919.054 | 1327.275 | 1122.334 | 1773.746 | 1099.463 | 848.215 | 624.101 |
| Surface | Non-saline | 1675 | 209.958 | 38.031 | 21.307 | 560.974 | 267.582 | 201.926 | 659.779 | 364.309 | 250.559 | 136.315 |
|  | Saline | 2956 | 609.448 | 242.789 | 155.392 | 1953.439 | 1542.279 | 1326.442 | 1516.418 | 1014.628 | 757.505 | 235.971 |
| Aquatic | Non-saline | 1218 | 306.709 | 123.823 | 80.030 | 507.457 | 312.123 | 237.334 | 394.588 | 235.160 | 173.751 | 65.414 |
|  | Saline | 1362 | 171.579 | 64.203 | 44.253 | 608.006 | 358.440 | 271.782 | 504.320 | 279.931 | 182.997 | 125.179 |
| Animal |  | 816 | 253.182 | 114.402 | 74.539 | 322.668 | 216.777 | 176.895 | 311.986 | 163.657 | 103.840 | 34.741 |
| Plant |  | 3527 | 329.969 | 23.389 | 11.400 | 1804.991 | 1440.221 | 1275.512 | 1811.139 | 1171.827 | 863.883 | 170.236 |
| Sediment |  | 4421 | 1613.380 | 543.674 | 241.657 | 2495.497 | 1794.819 | 1470.768 | 2215.265 | 1268.355 | 812.428 | 310.219 |
| Soil |  | 4955 | 1744.593 | 381.458 | 134.544 | 1140.601 | 770.768 | 664.830 | 1820.738 | 1101.519 | 828.295 | 295.596 |
| Surface |  | 1590 | 305.510 | 62.887 | 32.283 | 901.251 | 532.288 | 363.895 | 716.174 | 445.016 | 327.578 | 89.344 |
| Aquatic |  | 1549 | 292.213 | 101.645 | 65.219 | 573.845 | 325.476 | 244.807 | 625.627 | 402.172 | 309.768 | 62.424 |
| Non-saline |  | 3185 | 1116.316 | 300.138 | 135.690 | 974.957 | 587.149 | 478.888 | 1372.830 | 755.683 | 499.641 | 167.748 |
| Saline |  | 3209 | 509.192 | 135.442 | 80.469 | 1609.731 | 1161.427 | 965.205 | 1529.766 | 929.080 | 636.474 | 205.554 |
| Free-living |  | 2862 | 816.144 | 258.879 | 150.452 | 999.189 | 606.472 | 486.258 | 1245.129 | 746.131 | 550.564 | 145.757 |
| Host-associated |  | 1884 | 155.407 | 14.484 | 7.749 | 924.519 | 830.559 | 789.898 | 1011.591 | 740.404 | 575.552 | 130.291 |

**Table S9.** Results of permutation tests comparing estimated network heterogeneities of weighted networks representing 14 group × site combinations in the EMP dataset. Values shown are *p* values from permutation tests comparing network heterogeneities in Site 1 and Site 2 (estimates being compared are given in the first 14 rows of Table S8). See Appendix D for the details of the algorithm used for the permutation tests.

| Group | Site 1 | Site 2 | Network Heterogeneity in Node Abundances |  |  |  | Network Heterogeneity in Link Weights |  |  | Overall Network Heterogeneity |  |  |  |
| --- | --- | --- | --- | --- | --- | --- | --- | --- | --- | --- | --- | --- | --- |
| | | | $q=0$ | $q=1$ | $q=2$ | $q=3$ | $q=1$ | $q=2$ | $q=3$ | $q=1$ | $q=2$ | $q=3$ | $V/M$ |
| Animal | corpus | distal gut | 0 | 0 | 0 | 0 | 0 | 0 | 0 | 0 | 0 | 0 | 0.91 |
|  |  | proximal gut | 0 | 0 | 0 | 0 | 0 | 0 | 0 | 0 | 0 | 0.01 | 0.44 |
|  |  | secretion | 0 | 0 | 0 | 0 | 0.05 | 0.08 | 0.09 | 0 | 0 | 0 | 0.88 |
|  |  | surface | 0 | 0 | 0 | 0.03 | 0.48 | 0.85 | 0.86 | 0 | 0 | 0 | 0.84 |
|  | distal gut | proximal gut | 0 | 0 | 0.01 | 0.82 | 0.98 | 0.99 | 0.98 | 0.11 | 0.52 | 0.69 | 1.00 |
|  |  | secretion | 0 | 0 | 0 | 0 | 0 | 0 | 0 | 0 | 0.01 | 0.17 | 0.35 |
|  |  | surface | 0 | 0 | 0 | 0 | 0 | 0 | 0 | 0 | 0.46 | 0.93 | 0.69 |
|  | proximal gut | secretion | 0 | 0 | 0 | 0 | 0 | 0 | 0 | 0.24 | 0.7 | 0.86 | 0.98 |
|  |  | surface | 0 | 0.02 | 0 | 0 | 0.13 | 0.17 | 0.25 | 0.76 | 0.72 | 0.62 | 1.00 |
|  | secretion | surface | 0 | 0 | 0 | 0.15 | 0.1 | 0.9 | 0.87 | 0 | 0.01 | 0.03 | 0.30 |
| Plant | corpus | rhizosphere | 0 | 0 | 0 | 0 | 0 | 0 | 0 | 0 | 0 | 0 | 1.00 |
|  |  | surface | 0 | 0 | 0 | 0 | 0.18 | 0.45 | 0.6 | 0.78 | 0.7 | 0.6 | 0.99 |
|  | rhizosphere | surface | 0 | 0 | 0 | 0 | 0 | 0 | 0 | 0 | 0 | 0.02 | 0.72 |
| Sediment | non-saline | saline | 0 | 0.06 | 0.55 | 0.22 | 0.88 | 0.45 | 0.68 | 0.14 | 0.08 | 0.04 | 0.61 |
| Surface | non-saline | saline | 0.08 | 0 | 0 | 0 | 0 | 0 | 0 | 0.04 | 0.01 | 0 | 1.00 |
| Aquatic | non-saline | saline | 0 | 0 | 0 | 0 | 0.54 | 0.73 | 0.7 | 0.09 | 0.59 | 0.96 | 0.64 |

**Table S10.** Results of permutation tests comparing estimated network heterogeneities of weighted networks representing 10 groups in the EMP dataset. Values shown are *p* values from permutation tests comparing network heterogeneities in Group 1 and Group 2 (estimates being compared are given in the last 10 rows of Table S8). See Appendix D for the details of the algorithm used for the permutation tests.

| Group 1 | Group 2 | Network Heterogeneity in Node Abundances |  |  |  | Network Heterogeneity in Link Weights |  |  | Overall Network Heterogeneity |  |  |  |
| --- | --- | --- | --- | --- | --- | --- | --- | --- | --- | --- | --- | --- |
| | | $q=0$ | $q=1$ | $q=2$ | $q=3$ | $q=1$ | $q=2$ | $q=3$ | $q=1$ | $q=2$ | $q=3$ | $V/M$ |
| Animal | Plant | 0 | 0.36 | 0 | 0 | 0 | 0 | 0 | 0 | 0 | 0 | 0.24 |
|  | Sediment | 0 | 0 | 0 | 0 | 0 | 0 | 0 | 0 | 0 | 0 | 0.06 |
|  | Soil | 0 | 0 | 0 | 0.04 | 0 | 0 | 0 | 0 | 0 | 0 | 0.20 |
|  | Surface | 0 | 0.1 | 0 | 0 | 0 | 0.02 | 0.52 | 0 | 0 | 0 | 0.66 |
|  | Aquatic | 0 | 0.23 | 0.48 | 0.41 | 0 | 0.2 | 0.36 | 0 | 0 | 0 | 0.55 |
| Plant | Sediment | 0 | 0 | 0 | 0 | 0 | 0.06 | 0.28 | 0.01 | 0.62 | 0.79 | 0.41 |
|  | Soil | 0 | 0 | 0 | 0 | 0 | 0 | 0 | 0.95 | 0.72 | 0.89 | 0.23 |
|  | Surface | 0 | 0.78 | 0.02 | 0 | 0 | 0 | 0 | 0 | 0 | 0 | 0.49 |
|  | Aquatic | 0 | 0.56 | 0 | 0 | 0 | 0 | 0 | 0 | 0 | 0 | 0.33 |
| Sediment | Soil | 0 | 0.14 | 0.09 | 0.04 | 0 | 0 | 0 | 0.02 | 0.48 | 0.94 | 0.96 |
|  | Surface | 0 | 0 | 0 | 0 | 0 | 0 | 0 | 0 | 0 | 0.02 | 0.24 |
|  | Aquatic | 0 | 0 | 0 | 0 | 0 | 0 | 0 | 0 | 0 | 0 | 0.08 |
| Soil | Surface | 0 | 0 | 0 | 0 | 0.41 | 0.01 | 0 | 0 | 0 | 0 | 0.35 |
|  | Aquatic | 0 | 0 | 0 | 0 | 0 | 0 | 0 | 0 | 0 | 0 | 0.21 |
| Surface | Aquatic | 0 | 0.71 | 0.04 | 0 | 0.11 | 0.59 | 0.8 | 0.3 | 0.65 | 0.85 | 0.87 |
| Non-saline | Saline | 0 | 0 | 0 | 0.04 | 0.04 | 0.03 | 0.03 | 0.31 | 0.27 | 0.45 | 0.91 |
| Free-living | Host-associated | 0 | 0 | 0 | 0 | 0.09 | 0 | 0 | 0.13 | 0.96 | 0.8 | 0.94 |

**Table S11.** Parameter estimates for scaling heterogeneity scaling in the EMP dataset using Taylor’s Power Law of Networks (TPLoN). Groups and group × site combinations as in Table S1. Values given include sample size *N*, the parameters *a* and *b* of the TPLoN (*V = aM*^*b*^), the derived measure 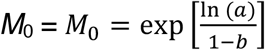, and measures of fit of the model to the data. See Appendix D for further discussion of the computation of these parameters.

| Group | Site | $b$ | $\ln(a)$ | $M_0$ | $R$ | $P$-value | $N$ |
| --- | --- | --- | --- | --- | --- | --- | --- |
| Animal | Corpus | 2.003 | 3.375 | 0.035 | 0.996 | <0.001 | 106 |
|  | Distal gut | 1.554 | 2.890 | 0.005 | 0.930 | <0.001 | 1696 |
|  | Proximal gut | 1.879 | 3.523 | 0.018 | 0.977 | <0.001 | 888 |
|  | Secretion | 1.812 | 3.684 | 0.011 | 0.981 | <0.001 | 631 |
|  | Surface | 1.883 | 4.664 | 0.005 | 0.984 | <0.001 | 1124 |
| Plant | Corpus | 1.697 | 2.084 | 0.050 | 0.972 | <0.001 | 334 |
|  | Rhizosphere | 1.658 | 2.190 | 0.036 | 0.933 | <0.001 | 4418 |
|  | Surface | 1.964 | 4.287 | 0.012 | 0.986 | <0.001 | 737 |
| Sediment | Non-saline | 1.756 | 2.870 | 0.022 | 0.926 | <0.001 | 2768 |
|  | Saline | 1.680 | 3.310 | 0.008 | 0.958 | <0.001 | 4345 |
| Surface | Non-saline | 1.688 | 3.674 | 0.005 | 0.962 | <0.001 | 1675 |
|  | Saline | 1.661 | 3.417 | 0.006 | 0.965 | <0.001 | 2956 |
| Aquatic | Non-saline | 1.836 | 4.343 | 0.006 | 0.980 | <0.001 | 1218 |
|  | Saline | 1.932 | 3.844 | 0.016 | 0.977 | <0.001 | 1362 |
| Animal |  | 1.719 | 3.443 | 0.008 | 0.980 | <0.001 | 816 |
| Plant |  | 1.506 | 2.795 | 0.004 | 0.966 | <0.001 | 3527 |
| Sediment |  | 1.601 | 3.654 | 0.002 | 0.938 | <0.001 | 4421 |
| Soil |  | 1.654 | 3.791 | 0.003 | 0.963 | <0.001 | 4955 |
| Surface |  | 1.731 | 3.958 | 0.004 | 0.965 | <0.001 | 1590 |
| Aquatic |  | 1.801 | 4.112 | 0.006 | 0.968 | <0.001 | 1549 |
| Non-saline |  | 1.550 | 3.748 | 0.001 | 0.947 | <0.001 | 3185 |
| Saline |  | 1.560 | 3.613 | 0.002 | 0.958 | <0.001 | 3209 |
| Free-living |  | 1.618 | 3.980 | 0.002 | 0.964 | <0.001 | 2862 |
| Host-associated |  | 1.462 | 2.919 | 0.002 | 0.985 | <0.001 | 1884 |

**Table S12.**
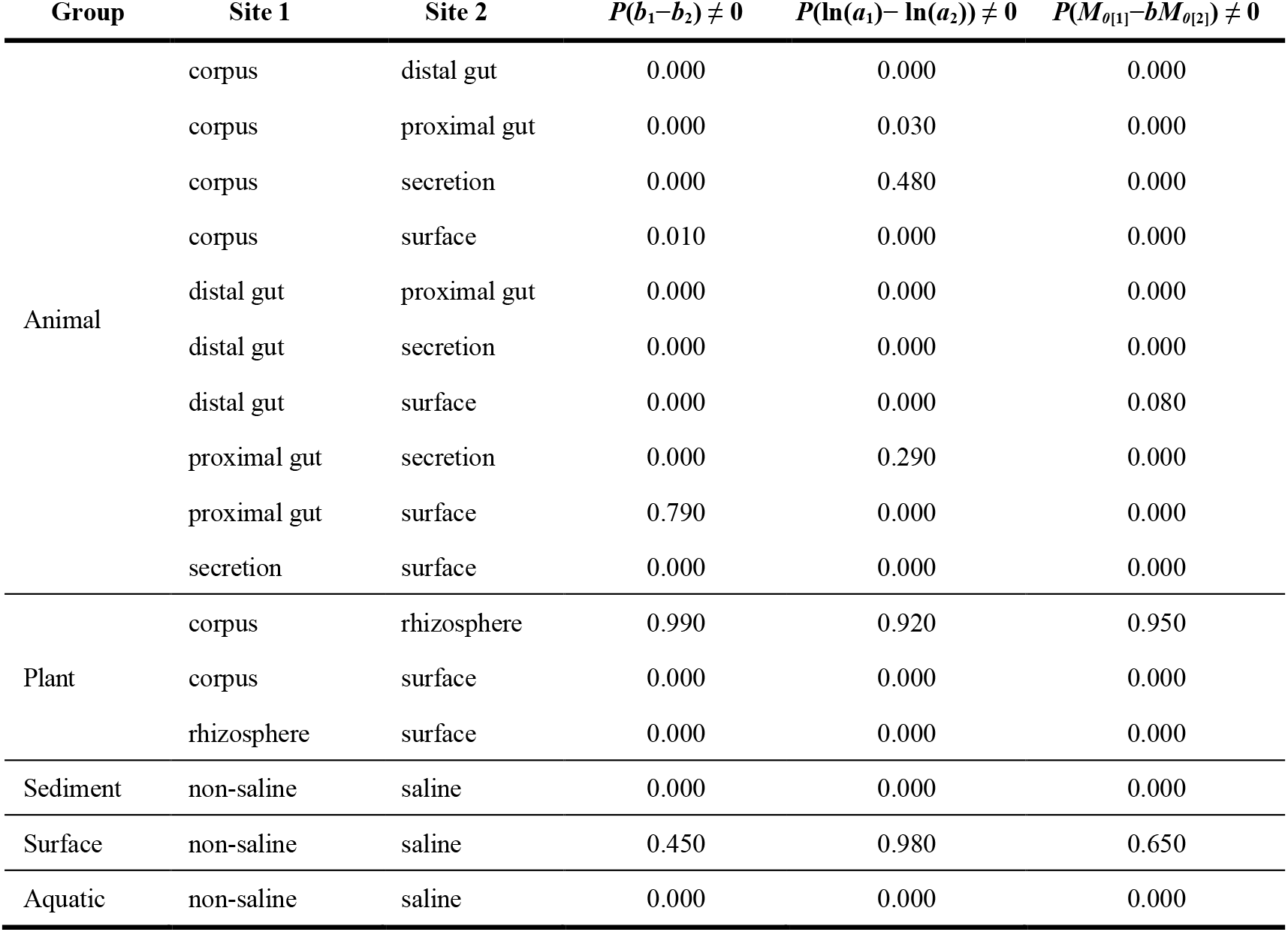
Results of permutation tests comparing TPLoN parameters of the weighted networks representing the 14 group × site combinations in the EMP dataset (Table S1). Values shown are *p* values from permutation tests on values of *b*, ln(*a*), and *M*_0_ given in the first 14 rows of Table S11.

**Table S13.** Results of permutation tests comparing TPLoN parameters of the weighted networks representing the 10 groups in the EMP dataset (Table S1). Values shown are *p* values from permutation tests on values of *b*, ln(*a*), and *M*_0_ given in the first 14 rows of Table S11.

| Group 1 | Group 2 | $P(b_1 - b_2) \neq 0$ | $P(\ln(a_1) - \ln(a_2)) \neq 0$ | $P(M_{\theta(1)} - bM_{\theta(2)}) \neq 0$ |
| --- | --- | --- | --- | --- |
| Animal | Plant | 0.000 | 0.000 | 0.000 |
|  | Sediment | 0.000 | 0.000 | 0.000 |
|  | Soil | 0.000 | 0.000 | 0.000 |
|  | Surface | 0.730 | 0.000 | 0.000 |
|  | Aquatic | 0.000 | 0.000 | 0.000 |
| Plant | Sediment | 0.000 | 0.000 | 0.000 |
|  | Soil | 0.000 | 0.000 | 0.000 |
|  | Surface | 0.000 | 0.000 | 0.000 |
|  | Aquatic | 0.000 | 0.000 | 0.000 |
| Sediment | Soil | 0.000 | 0.000 | 0.000 |
|  | Surface | 0.000 | 0.000 | 0.000 |
|  | Aquatic | 0.000 | 0.000 | 0.000 |
| Soil | Surface | 0.000 | 0.000 | 0.000 |
|  | Aquatic | 0.000 | 0.000 | 0.000 |
| Surface | Aquatic | 0.000 | 0.320 | 0.000 |
| Non-saline | Saline | 0.630 | 0.000 | 0.000 |
| Free-living | Host-associated | 0.000 | 0.000 | 0.020 |

**Table S14.** Details of the nine switches in heterogeneity (*H*_*i*_) types of a given OTU between two networks in the 14 group × site combinations of the EMP dataset (see Tables 2 and 3 in main text for overall summaries and Table S1 for a summary of the EMP dataset). See Dataset S3 for the full data and Table S16 for its associated metadata.

| Group | Site 1 | Site 2 | $H \rightarrow H$ | $H \rightarrow R$ | $H \rightarrow U$ | $R \rightarrow H$ | $R \rightarrow R$ | $R \rightarrow U$ | $U \rightarrow H$ | $U \rightarrow R$ | $U \rightarrow U$ | Total |
| --- | --- | --- | --- | --- | --- | --- | --- | --- | --- | --- | --- | --- |
| **For OTUs in Site 1 whose $H_i > H_i$ in site 2** |  |  |  |  |  |  |  |  |  |  |  |  |
| Animal | corpus | distal gut | 0 | 0 | 0 | 0 | 0 | 0 | 0 | 0 | 0 | 0 |
|  | corpus | proximal gut | 0 | 0 | 0 | 0 | 0 | 0 | 0 | 0 | 0 | 0 |
|  | corpus | secretion | 0 | 1 | 6 | 0 | 0 | 6 | 0 | 0 | 0 | 13 |
|  | corpus | surface | 0 | 2 | 2 | 0 | 2 | 4 | 0 | 0 | 0 | 10 |
|  | distal gut | proximal gut | 0 | 4 | 5 | 0 | 0 | 0 | 0 | 0 | 0 | 9 |
|  | distal gut | secretion | 1 | 0 | 1 | 0 | 0 | 0 | 0 | 0 | 0 | 2 |
|  | distal gut | surface | 1 | 10 | 4 | 0 | 0 | 0 | 0 | 0 | 0 | 15 |
|  | proximal gut | secretion | 0 | 4 | 9 | 0 | 0 | 0 | 0 | 0 | 0 | 13 |
|  | proximal gut | surface | 2 | 8 | 3 | 0 | 0 | 1 | 0 | 0 | 0 | 14 |
|  | secretion | surface | 10 | 18 | 8 | 0 | 6 | 3 | 0 | 0 | 1 | 46 |
| Plant | corpus | rhizosphere | 8 | 5 | 4 | 0 | 0 | 1 | 0 | 0 | 0 | 18 |
|  | corpus | surface | 0 | 0 | 0 | 0 | 0 | 0 | 0 | 0 | 0 | 0 |
|  | rhizosphere | surface | 1 | 1 | 1 | 0 | 0 | 0 | 0 | 0 | 0 | 3 |
| Sediment | non-saline | saline | 8 | 2 | 0 | 0 | 0 | 0 | 0 | 0 | 0 | 10 |
| Surface | non-saline | saline | 0 | 0 | 0 | 0 | 0 | 1 | 0 | 0 | 0 | 1 |
| Aquatic | non-saline | saline | 0 | 0 | 1 | 0 | 0 | 1 | 0 | 0 | 0 | 2 |
| **For OTUs in Site 1 whose $H_i < H_i$ in site 2** |  |  |  |  |  |  |  |  |  |  |  |  |
| Animal | corpus | distal gut | 0 | 0 | 0 | 0 | 0 | 0 | 1 | 0 | 0 | 1 |
|  | corpus | proximal gut | 0 | 0 | 0 | 0 | 0 | 0 | 0 | 1 | 0 | 1 |
|  | corpus | secretion | 1 | 0 | 0 | 0 | 0 | 1 | 1 | 0 | 0 | 3 |
|  | corpus | surface | 1 | 0 | 0 | 0 | 0 | 1 | 1 | 0 | 0 | 3 |
|  | distal gut | proximal gut | 7 | 0 | 0 | 10 | 0 | 0 | 3 | 3 | 0 | 23 |
|  | distal gut | secretion | 4 | 0 | 0 | 19 | 1 | 0 | 4 | 3 | 0 | 31 |
|  | distal gut | surface | 5 | 0 | 0 | 16 | 3 | 0 | 5 | 2 | 0 | 31 |
|  | proximal gut | secretion | 0 | 0 | 0 | 0 | 0 | 0 | 3 | 1 | 0 | 4 |
|  | proximal gut | surface | 1 | 0 | 0 | 1 | 0 | 0 | 4 | 1 | 0 | 7 |
|  | secretion | surface | 25 | 1 | 0 | 19 | 2 | 0 | 17 | 21 | 6 | 91 |
| Plant | corpus | rhizosphere | 0 | 0 | 0 | 0 | 0 | 1 | 1 | 1 | 1 | 4 |
|  | corpus | surface | 0 | 0 | 0 | 0 | 0 | 0 | 0 | 0 | 0 | 0 |
|  | rhizosphere | surface | 0 | 0 | 0 | 0 | 0 | 0 | 0 | 0 | 0 | 0 |
| Sediment | non-saline | saline | 169 | 0 | 0 | 41 | 0 | 0 | 9 | 0 | 0 | 219 |
| Surface | non-saline | saline | 36 | 0 | 0 | 25 | 0 | 0 | 7 | 0 | 0 | 68 |
| Aquatic | non-saline | saline | 2 | 0 | 0 | 21 | 0 | 0 | 17 | 0 | 0 | 40 |

**Table S15.** Details of the nine switches in heterogeneity (*H*_*i*_) types of a given OTU between two networks in the 10 groups of the EMP dataset (see Tables 2 and 3 in main text for overall summaries and Table S1 for a summary of the EMP dataset). See Dataset S4 for the full data and Table S17 for its associated metadata.

| Group 1 | Group 2 | $H \rightarrow H$ | $H \rightarrow R$ | $H \rightarrow U$ | $R \rightarrow H$ | $R \rightarrow R$ | $R \rightarrow U$ | $U \rightarrow H$ | $U \rightarrow R$ | $U \rightarrow U$ | Total |
| --- | --- | --- | --- | --- | --- | --- | --- | --- | --- | --- | --- |
| **For OTUs in Group 1 whose $H_i > H_i$ in Group 2** |  |  |  |  |  |  |  |  |  |  |  |
| Animal | Plant | 0 | 0 | 1 | 0 | 0 | 1 | 0 | 0 | 0 | 2 |
|  | Sediment | 0 | 0 | 0 | 0 | 0 | 0 | 0 | 0 | 0 | 0 |
|  | Soil | 0 | 4 | 1 | 0 | 0 | 1 | 0 | 0 | 0 | 6 |
|  | Surface | 2 | 0 | 1 | 0 | 0 | 0 | 0 | 0 | 1 | 4 |
|  | Aquatic | 0 | 0 | 1 | 0 | 0 | 0 | 0 | 0 | 0 | 1 |
| Plant | Sediment | 171 | 5 | 0 | 0 | 0 | 0 | 0 | 0 | 0 | 176 |
|  | Soil | 575 | 175 | 21 | 0 | 1 | 1 | 0 | 0 | 1 | 774 |
|  | Surface | 20 | 9 | 1 | 0 | 0 | 2 | 0 | 0 | 0 | 32 |
|  | Aquatic | 26 | 19 | 7 | 0 | 1 | 1 | 0 | 0 | 0 | 54 |
| Sediment | Soil | 232 | 95 | 14 | 0 | 0 | 0 | 0 | 0 | 0 | 341 |
|  | Surface | 36 | 12 | 0 | 0 | 0 | 0 | 0 | 0 | 0 | 48 |
|  | Aquatic | 42 | 39 | 16 | 0 | 0 | 2 | 0 | 0 | 0 | 99 |
| Soil | Surface | 17 | 10 | 2 | 1 | 0 | 0 | 0 | 0 | 0 | 30 |
|  | Aquatic | 17 | 19 | 7 | 0 | 2 | 1 | 0 | 0 | 0 | 46 |
| Surface | Aquatic | 13 | 28 | 23 | 0 | 0 | 4 | 0 | 0 | 0 | 68 |
| Non-saline | Saline | 1 | 0 | 2 | 0 | 0 | 1 | 0 | 0 | 0 | 4 |
| Free-living | Host-associated | 30 | 12 | 7 | 0 | 2 | 1 | 0 | 0 | 0 | 52 |
| **For OTUs in Group 1 whose $H_i < H_i$ in Group 2** |  |  |  |  |  |  |  |  |  |  |  |
| Animal | Plant | 1 | 0 | 0 | 9 | 1 | 0 | 7 | 1 | 0 | 19 |
|  | Sediment | 0 | 0 | 0 | 5 | 0 | 0 | 1 | 0 | 0 | 6 |
|  | Soil | 5 | 1 | 0 | 11 | 0 | 0 | 3 | 2 | 0 | 22 |
|  | Surface | 14 | 0 | 0 | 27 | 1 | 0 | 16 | 3 | 1 | 62 |
|  | Aquatic | 3 | 0 | 0 | 5 | 0 | 0 | 5 | 1 | 0 | 14 |
| Plant | Sediment | 198 | 0 | 0 | 20 | 0 | 0 | 6 | 0 | 0 | 224 |
|  | Soil | 616 | 2 | 0 | 8 | 2 | 0 | 6 | 1 | 0 | 635 |
|  | Surface | 54 | 1 | 0 | 11 | 1 | 0 | 11 | 2 | 0 | 80 |
|  | Aquatic | 12 | 0 | 0 | 7 | 0 | 0 | 6 | 0 | 0 | 25 |
| Sediment | Soil | 170 | 0 | 0 | 2 | 0 | 0 | 0 | 0 | 0 | 172 |
|  | Surface | 41 | 0 | 0 | 4 | 0 | 0 | 1 | 0 | 0 | 46 |
|  | Aquatic | 41 | 0 | 0 | 4 | 0 | 0 | 0 | 0 | 0 | 45 |
| Soil | Surface | 66 | 1 | 0 | 51 | 0 | 0 | 19 | 0 | 0 | 137 |
|  | Aquatic | 16 | 0 | 0 | 13 | 0 | 0 | 7 | 0 | 0 | 36 |
| Surface | Aquatic | 7 | 0 | 0 | 3 | 0 | 0 | 2 | 0 | 0 | 12 |
| Non-saline | Saline | 43 | 0 | 0 | 15 | 0 | 0 | 6 | 0 | 0 | 64 |
| Free-living | Host-associated | 148 | 0 | 0 | 25 | 2 | 1 | 1 | 1 | 0 | 178 |

**Table S16.** Metadata for Dataset S3. **Filename**: DatasetS3.csv **Format**: comma-separated values (.csv) **Description**: Comparisons of node heterogeneities between OTUs that co-occur in all possible pairs of the 14 group × site combinations in the EMP dataset (Table S1). **Variable definitions**:

| Column | Variable | Type and description |
| --- | --- | --- |
| 1 | EMP_group | Character: Group assignment for each site in columns 2 and 3 (see column 1 of Table S1) |
| 2 | EMP_site_1 | Character: Site assignment for network 1 (see column 2 of Table S2) |
| 3 | EMP_site_2 | Character: Site assignment for network 2 (see column 2 of Table S2) |
| 3 | OTU | Character: Operational Taxonomic Unit.<br>Formatted as:<br><level>_<name>_<number>, where <level> is one of s (species), g (genus), f (family), o (order), c (class), or p (phylum); <name> is either a taxonomic name or alphanumeric identifier for undescribed taxa; and <number> is a reference number for the taxon. |
| 4 | Hi_1 | Real: Node heterogeneity of the OTU in the group × site 1 combination |
| 5 | Hi_2 | Real: Node heterogeneity of the OTU in the group × site 2 combination |
| 6 | Hi_1_gt_Hi_2 | Logical (True/False): Is Hi_1 > Hi_2? |
| 7 | Delta_Hi | Real: Hi_1 – Hi_2 |
| 8 | P_value | Real: P-value of Delta_Hi from a permutation test |
| 6 | H_type_1 | Character: one of “heterogeneous”, “uniform”, or “random”. This variable is the assignment to heterogeneous ($H_i > 1$), uniform ($H_i < 1$), or random (neither heterogeneous or uniform) cluster in the weighted network given by the group × site_1 combination. |
| 6 | H_type_2 | Character: one of “heterogeneous”, “uniform”, or “random”. This variable is the assignment to heterogeneous ($H_i > 1$), uniform ($H_i < 1$), or random (neither heterogeneous or uniform) cluster in the weighted network given by the group × site_2 combination. |

**Table S17.** Metadata for Dataset 4. **Filename**: DatasetS3.csv **Format**: comma-separated values (.csv) **Description**: Comparisons of node heterogeneities between OTUs that co-occur in all possible pairs of the 10 group combinations in the EMP dataset (Table S1). **Variable definitions**:

| Column | Variable | Type and description |
| --- | --- | --- |
| 1 | EMP_group_1 | Character: Group assignment for network 1 (see column 1 of Table S1) |
| 2 | EMP_group_2 | Character: Group assignment for network 2 (see column 1 of Table S1) |
| 3 | OTU | Character: Operational Taxonomic Unit.<br>Formatted as:<br><level>_<name>_<number>, where <level> is one of s (species), g (genus), f (family), o (order), c (class), or p (phylum); <name> is either a taxonomic name or alphanumeric identifier for undescribed taxa; and <number> is a reference number for the taxon. |
| 4 | Hi_1 | Real: Node heterogeneity of the OTU in the group × site 1 combination |
| 5 | Hi_2 | Real: Node heterogeneity of the OTU in the group × site 2 combination |
| 6 | Hi_1_gt_Hi_2 | Logical (True/False): Is $H_{i\_1} > H_{i\_2}$? |
| 7 | Delta_Hi | Real: $H_{i\_1} - H_{i\_2}$ |
| 8 | P_value | Real: P-value of Delta_Hi from a permutation test |
| 6 | H_type_1 | Character: one of “heterogeneous”, “uniform”, or “random”. This variable is the assignment to heterogeneous ($H_i > 1$), uniform ($H_i < 1$), or random (neither heterogeneous or uniform) cluster in the weighted network given by the group × site_1 combination. |
| 6 | H_type_2 | Character: one of “heterogeneous”, “uniform”, or “random”. This variable is the assignment to heterogeneous ($H_i > 1$), uniform ($H_i < 1$), or random (neither heterogeneous or uniform) cluster in the weighted network given by the group × site_2 combination. |

